# Evolutionary genomics of a natural *Wolbachia* superinfection in the dengue mosquito *Aedes albopictus*

**DOI:** 10.64898/2026.06.26.734713

**Authors:** Thomas L Schmidt, Julien Martinez, Jonathan P Day, Steven P. Sinkins, John J Welch, Francis M Jiggins

## Abstract

*Aedes albopictus*, an invasive mosquito vector of human pathogens, is naturally infected by two strains of the endosymbiotic bacterium *Wolbachia, w*AlbA and *w*AlbB. Here, we present a comprehensive genomic analysis of these *Wolbachia* strains, their associated plasmids, and host mitochondrial DNA (mtDNA). We observed extensive variation in *Wolbachia* densities—spanning over three orders of magnitude—indicating that infections often exist at extremely low levels and this variation is largely driven by environmental factors. Phylogenetic analyses revealed concordant evolutionary histories among mitochondrial, *Wolbachia*, and plasmid genomes, with no strong evidence for horizontal or paternal transmission. We estimate that the common ancestor of the superinfection existed ∼6,000—9,000 years ago. Patterns of genetic structure and divergence indicate that the subsequent geographical spread of *Wolbachia* has mirrored host population history. Comparative rate analyses revealed that *w*AlbA evolves more slowly than *w*AlbB, consistent with a reduced mutation rate. Despite the close relationship between wAlbB lineages, gene content in this strain is highly variable, with 184 genes exhibiting presence-absence or copy number variation across strains, including genes involved in inducing cytoplasmic incompatibility. Several of these genes reside in repetitive or prophage regions, indicating that transposable elements appear to drive structural genomic variation in *Wolbachia* populations. Together, our results highlight extensive natural diversity within *Wolbachia* populations and provide a foundation for understanding symbiont dynamics in nature and to inform the selection of strains for *Wolbachia*-based vector control programs.

## Introduction

*Wolbachia* is a bacterium that inhabits the cells of invertebrate hosts and is found in about half of all terrestrial arthropod species (Weinert et al. 2015; Bailly-Bechet et al. 2017). Like mitochondria, *Wolbachia* is transmitted intergenerationally via the female germline, which can lead to similar structure in mitochondrial and *Wolbachia* phylogenies (Richardson et al. 2012). However, over longer evolutionary timescales horizontal transmission occurs between species, as the phylogenies of insect hosts and their *Wolbachia* are rarely congruent (O’Neill et al. 1992; Turelli et al. 2022). Analyses of *Wolbachia* phylogenies have revealed that *Wolbachia-*host associations can be dynamic, with a lineage of *Wolbachia* spreading across eight *Drosophila* species in ∼14,000 years. While horizontal transmission is a rare event that shapes the *Wolbachia-*host associations over the long term, failed vertical transmission, with infected females producing uninfected offspring, is common in many populations (Ahmed et al. 2015; Li et al. 2017; Schmidt et al. 2018). This can result in uninfected and infected individuals stably coexisting in the same population (Fine 1978).

*Wolbachia* can induce a range of reproductive phenotypes in hosts that aid its spread through populations. The most common is cytoplasmic incompatibility, in which the eggs of females without *Wolbachia* will be sterile if mated with *Wolbachia*-infected males, giving a fitness advantage to infected females (Hoffmann and Turelli 1997). Cytoplasmic incompatibility has been exploited for successful management of pests and disease vectors such as *Aedes* mosquitoes following transinfection of *Wolbachia* from other hosts (Hoffmann et al. 2011; Nazni et al. 2019; Zheng et al. 2019; Crawford et al. 2020; Beebe et al. 2021; Utarini et al. 2021). Aside from reproductive manipulations, *Wolbachia* infection can affect host fitness in numerous ways and these can show strong variation between (Yeap et al. 2011; Ross et al. 2017) and within (Ross et al. 2023) *Wolbachia* strains. These fitness effects can determine whether *Wolbachia* invade insect populations, making it of critical importance to select appropriate strains for control programs (Salje and Jiggins 2024).

*Wolbachia* is divided into supergroups, with supergroups A and B being the most common in insects (Werren, Zhang, et al. 1995). Hosts can be superinfected with two or more strains, with superinfections often covering both supergroups (Werren, Windsor, et al. 1995; Dobson et al. 2004; Ellegaard et al. 2013). Provided crosses between superinfected males and singly infected females are incompatible, superinfections can spread through singly-infected populations just as single infections can through uninfected populations, leading to selective sweeps of *Wolbachia* and coinherited mtDNA in host populations (Hurst and Jiggins 2005). Natural superinfections can exist stably across a species range, such as in the mosquito *Ae. albopictus,* where superinfected hosts have similar fitness to singly-infected hosts but with the added benefit of cytoplasmic incompatibility (Dobson et al. 2004). Despite its global distribution, *Ae. albopictus* shows considerable variability in strain density among individuals and populations as well as between sexes (Yang et al. 2022).

The ubiquity of *Wolbachia* and the strong phenotypic effects it confers on its hosts make it an important model of symbiosis and long-term coevolution. There are a small number of cases where single infections have been studied using population genomic methods (Richardson et al. 2012; Turelli et al. 2018; Cooper et al. 2019). These approaches can provide insights into the mechanisms giving rise to genetic variation in populations and variation in gene copy number and repeat content that may affect phenotype.

Phylogenetic comparisons of symbiont and mitochondrial genomes allow past patterns of horizontal and vertical transmission to be reconstructed. Previous genomic studies have not encompassed multiple, geographically distinct populations where host nuclear background and external environment vary. This can indicate where and when the infection originated, whether it spread passively with host migration or independently of host population history, and whether key traits like symbiont density are shaped by local environmental conditions or symbiont genetics. Previous genomic studies have also not considered superinfections, where the shared evolutionary history and environment of strains superinfecting the same host allow comparisons of evolutionary patterns and rates. Superinfections are of particular interest as the strains may be in resource competition with each other within the host. Although resource competition can reduce total *Wolbachia* density (Dutton and Sinkins 2004), it has been suggested that superinfections across A and B supergroups may be ecologically specialised and thus avoid competition through niche partitioning (Ellegaard et al. 2013).

We investigated the evolution of the *Wolbachia* superinfection found across globally distributed populations of *Ae. albopictus* (the Asian tiger mosquito). *Aedes albopictus* is a highly invasive disease vector native to Asia, that has undergone rapid and ongoing range expansion in recent decades, now occupying extensive regions spanning temperate to tropical climates worldwide. Globally it is the second most important vector of dengue after *Ae. aegypti* and is the primary vector in locations such as southern China (Yiguan et al. 2017). *Aedes albopictus* is superinfected with two *Wolbachia* strains from the A and B supergroups—*w*AlbA and *w*AlbB—and two plasmids associated with *w*AlbA (Martinez, Ant, et al. 2022). *w*AlbB is currently used in *Ae. aegypti* control programs as it strongly reduces arbovirus transmission in transinfected mosquitoes (Bian et al. 2010; Chouin-Carneiro et al. 2020). Accordingly, variation in *w*AlbB is of major public health interest (Ross et al. 2023). Most *Ae. albopictus* carry both *w*AlbA and *w*AlbB, though their densities are lower than in transinfected *Ae. aegypti* (Ant et al. 2018; Ant and Sinkins 2018), and qPCR analysis showed high density variation in both strains among individuals and populations including occasional loss of strains (Yang et al. 2022). *w*AlbA is typically found at lower densities than *w*AlbB (Dutton and Sinkins 2004). Although natural infections of wAlbA and wAlbB in *Ae. albopictus* are less effective at blocking dengue than when transinfected into *Ae. aegypti* (Lu et al. 2012), there is nevertheless some evidence that they reduce viral density in *Ae. albopictus* salivary glands (Mousson et al. 2012), pointing to a potential role in virus blocking (see also (Raquin et al. 2015)). If weaker virus blocking in *Ae. albopictus* is due to relatively lower *Wolbachia* densities, then knowledge of the environmental and genetic factors influencing density may be important for understanding disease risk.

Here we analyse *Wolbachia* and host genomes from across the range of *Ae. albopictus.* We find *Wolbachia* strains are detectable at far lower densities than when assessed with qPCR, showing extreme variation in density levels across individuals. *Aedes albopictus* mtDNA, *w*AlbA, *w*AlbB, and two plasmids have had perfect vertical transmission and spread ∼6,000—9,000 years ago, likely during an initial expansion of *Ae. albopictus* from Southeast Asia. However, rates of evolution differ; *w*AlbA evolves more slowly than *w*AlbB likely as a result of a reduced mutation rate. Finally, we find highly variable gene content in *w*AlbB across populations, including genes involved in inducing cytoplasmic incompatibility.

## 2. Results

### Genome sequences of *Wolbachia*, plasmids, and mitochondria from a global collection of *Ae. albopictus*

We selected 94 *Ae. albopictus* mosquitoes from a global set of samples that had been previously screened for *Wolbachia* infection using qPCR (Yang et al. 2022) and investigated genomically using ddRADseq (Schmidt et al. 2020; Schmidt et al. 2021). They came from sites in the presumed Asian native range as well as invasive range samples from Fiji (invaded ∼1988, Kay et al. (1995)), northeastern Papua New Guinea (Madang, invaded ∼1972 (Cooper et al. 1994)), Vanuatu (invaded ∼2011 (Guillaumot et al. 2012)), and the Torres Strait Islands in northern Australia (invaded 2004 (Ritchie et al. 2006)) (Figure 1). Samples were selected based on qPCR results to include mosquitoes with both *Wolbachia* strains (*n*=85) and those with only *w*AlbA (*n*=4, ‘A’ suffixes) or only *w*AlbB (*n*=5, ‘B’ suffixes), but within each subgroup selections were made randomly with respect to *Wolbachia* density. Samples were selected without regard to sex, though most (79) were female, 8 were male, and 7 of unknown sex.

**Figure 1.**
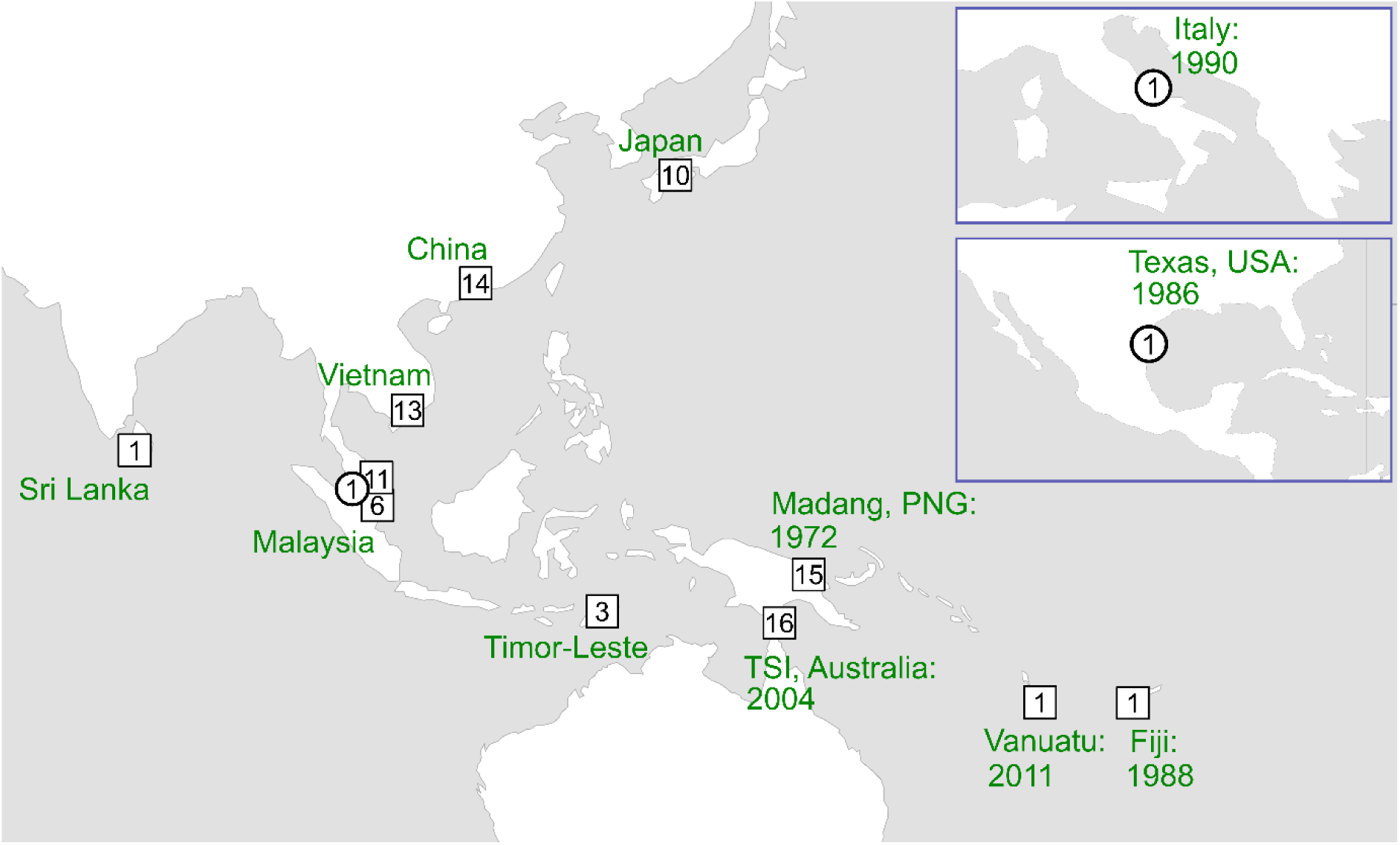
Sample locations and invasion dates. Numbers within shapes indicate sample sizes used in phylogenetic analyses. Text details known invasion times. TSI = Torres Strait Islands; PNG = Papua New Guinea. Squares show new genomes, circles published genomes.

We constructed Illumina sequencing libraries from each mosquito and used hybridisation probes to enrich for five genomes: the mtDNA genome, the *w*AlbA genome, the *w*AlbB genome, and two plasmid genomes reported from *w*AlbA (pWALBA1 and pWALBA2) (Martinez, Ant, et al. 2022). These data were combined with three previously sequenced strains, from Kuala Lumpur (*w*AlbA-JF2017: mtDNA and *w*AlbA), Texas, USA (*w*AlbB-Hou: mtDNA and wAlbB, invaded ∼1986 (Moore and Mitchell 1997)), and Italy (*w*AlbB-Uju: *w*AlbB, invaded ∼1990 (Dalla Pozza and Majori 1992)). Note that while *w*AlbB-Uju was introgressed into an Indonesian host background, the *Wolbachia* strain was sourced from Italy (Blagrove et al. 2012).

We used stringent criteria to ensure reads were mapped to the correct *Wolbachia* genome and subsequently discarded three low quality samples. For phylogenetic analyses, we required a minimum depth of 15X to call a nucleotide site and omitted individual genomes with >20% missing genotypes. This left 93 mtDNA genomes, 63 *w*AlbA genomes, 71 *w*AlbB genomes, 72 pWALBA1 genomes and 70 pWALBA2 genomes (Table S1). All five genomes were obtained from 59 samples (Table S1).

### *Wolbachia* can infect hosts at extremely low densities

There was considerable variation in the density of *Wolbachia* within mosquitoes estimated from sequencing depth, with the ratio of *Wolbachia* to mtDNA read depth varying by >10,000-fold for both *w*AlbA and *w*AlbB (Figure 2A). The most highly infected samples had ∼0.43 *Wolbachia* genomes for every mtDNA genome. In contrast to similar work on *Drosophila* (Richardson et al. 2012), there was no clear division of infected and uninfected samples. Instead, there was a continuum from the highest density samples down to a single sample that did not yield any reads mapping to the *w*AlbB genome (Figure 2A). *w*AlbA and *w*AlbB sequencing depths across individuals were positively correlated after accounting for variation in mitochondrial sequencing depth, as assessed using Standardized Major Axis (SMA) regression (see methods, Figure 2B). The *Wolbachia* density of these samples has been previously estimated by qPCR (Yang et al. 2022), and for both strains these results were positively but imperfectly correlated with our estimates (Figure S1; *w*AlbA *rho* = 0.5, *P* < 0.0001; *w*AlbB *rho* = 0.57, *P* < 0.0001). All nine samples classed as singly-infected by qPCR had reads that mapped to both genomes.

**Figure 2.**
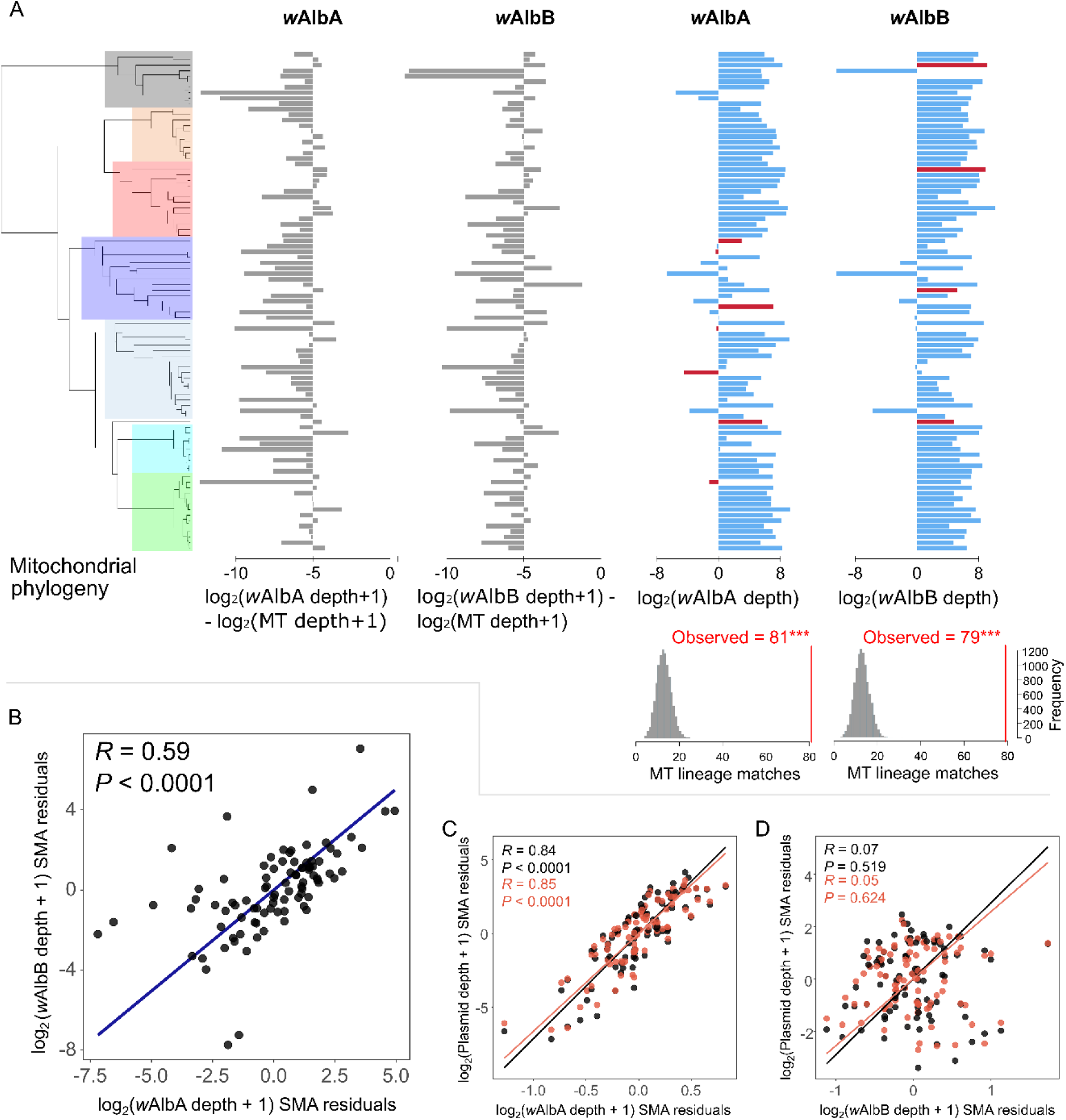
*Wolbachia* density and plasmid depth of coverage. (A) Left: Mitochondrial phylogeny with colors showing different lineages. Right: Barplots of relative density (gray) and sequencing depth (blue/red) for each *Wolbachia* genome per sample. Blue and red bars respectively indicate a match or a mismatch in mitochondrial lineage between each sample and its closest relative, based on Euclidean distances in the k-mer analysis. Bottom right: Distribution of the number of mitochondrial lineage matches drawn from 10,000 random permutations. P-value reflects the proportion of times the number of permuted matches was greater than or equal to the observed number of matches (red line). (B) Relationship between *w*AlbA and *w*AlbB sequencing depth after removing variation explained by mitochondrial sequencing depth. Points show residual sequencing depths, blue line represents the standardized major axis (SMA) regression. (C-D) Partial associations between plasmids and *Wolbachia* sequencing depths. Points show pWALBA1 (black) and pWALBA2 (orange) residual sequencing depths against wAlbA (C) or wAlbB (D) residual sequencing depth, after removing variation explained by mitochondrial depth and by the other Wolbachia strain. Lines represent SMA regressions per plasmid.

Sequence-based detection of *Wolbachia* is known to be more sensitive than qPCR (Pascar et al. 2023), but very low read counts could arise from sample contamination, ‘index hopping’ during sequencing or bioinformatic error. To test if samples with low *Wolbachia* sequencing depth were genuine infections, we analyzed the distribution of 20 bp *k*-mers overlapping SNPs in the *w*AlbA and *w*AlbB reference genomes. This *k*-mer approach was used because samples with extremely low *Wolbachia* depth were often missing >80% of genotype information, precluding a reliable inference of phylogenetic relatedness based on whole-genome variation. Using the raw sequencing reads, *k*-mers matching the reference and alternate alleles at each variable position in the *Wolbachia k*-mer database were counted and the ratio of these counts were used to calculate Euclidean distances between samples (see Materials and Methods). The closest relative of every sample was then identified and their respective mitochondrial lineages compared.

Under the assumption that *Wolbachia* is only transmitted maternally, we reasoned that target samples and their closest relative (based on *Wolbachia k*-mers) should belong to the same lineage in the mitochondrial phylogeny (Figure 2A; seven lineages highlighted). The observed number of mitochondrial lineage matches between samples and their closest neighbour was higher than expected by chance (10,000 random permutations; *P* < 0.0001 for both *w*AlbA and *w*AlbB), showing that our *Wolbachia k*-mer analysis predicts mitochondrial lineages (Figure 2A). However, the difference between the rate at which *Wolbachia* genomes <1X and >1X coverage were assigned to the correct mitochondrial lineage differed, with more mismatches for low coverage genomes (Fisher’s Exact Test: *p*=0.025). Nevertheless, among *Wolbachia* genomes with <1X coverage, all 9 *w*AlbB genomes and 8 of 12 *w*AlbA genomes were associated with their expected mitochondrial lineage. This included the samples with extremely low coverage, suggesting a real *Wolbachia* infection (Figure 2A).

### *Wolbachia* density is influenced by environment and geography

*w*AlbA and *w*AlbB densities could be influenced by *Wolbachia* genetics, the abiotic environment (e.g. temperature or environmental antibiotics) and the internal environment of the mosquito (e.g. larval crowding, nutrition (Dutton and Sinkins 2004), age (Ross et al. 2023) or mosquito genetics). All these factors may vary geographically. Furthermore, because the two symbionts share their environment and genetic history, they could all generate the correlation seen between *w*AlbA and *w*AlbB densities (Figure 2B). If *Wolbachia* density is controlled genetically, then we would expect related symbiont strains to have similar densities. Initially, we investigated this by including the phylogenetic subclade as a predictor in a linear model (the subclades were defined using the mitochondrial tree, allowing the inclusion of low coverage *Wolbachia* strains; Figure 3A). The model also included population of origin, sex and whether adults or larvae were sequenced.

**Figure 3.**
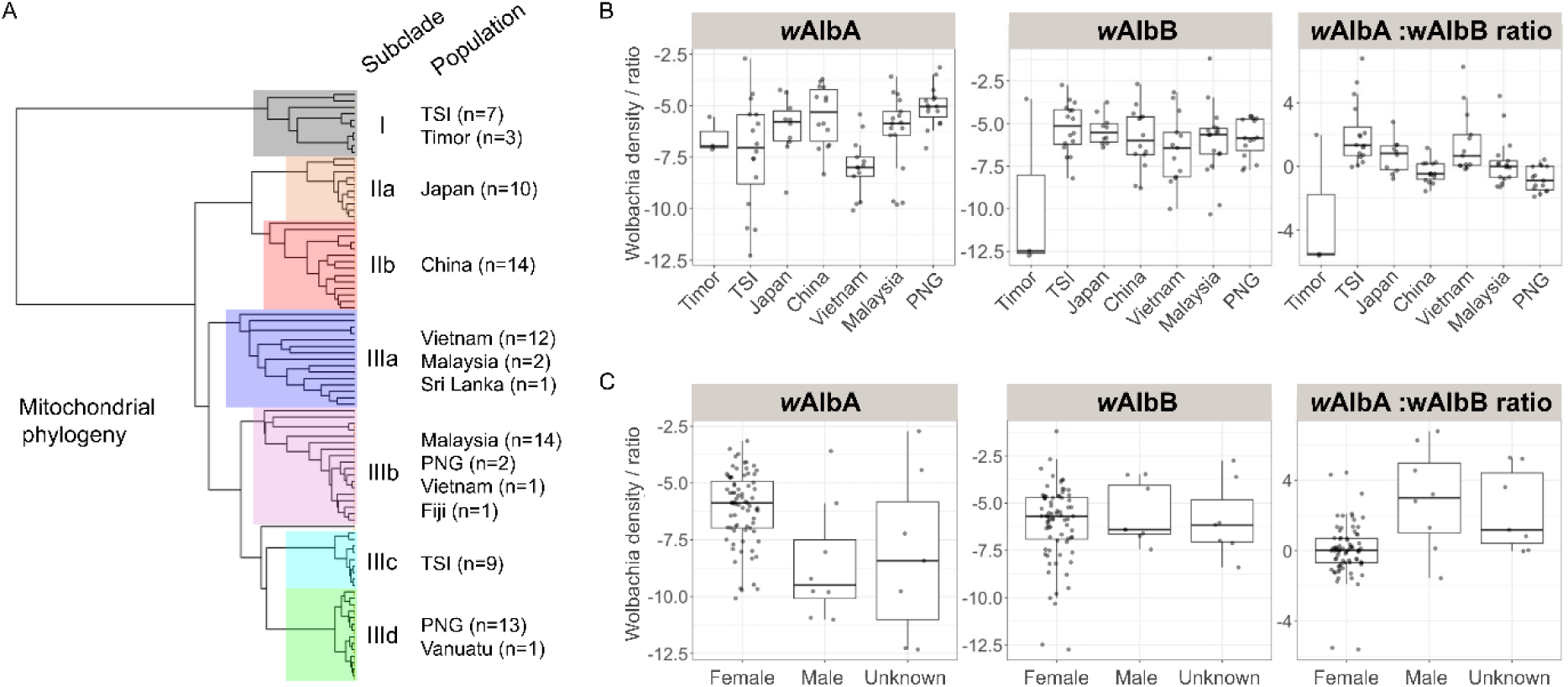
Predictors of variation in *Wolbachia* density. (A) Mitochondrial phylogeny indicating subclades and corresponding populations of origin of the samples in a given subclade (n: number of samples). (B) Variation in densities of *w*AlbA and *w*AlbB and in their ratio between populations. (C) Density variation between sexes. TSI = Torres Strait Islands; PNG = Papua New Guinea.

We found no evidence of genetic differences between subclades affecting the density of *w*AlbA or *w*AlbB (Table 1). We reasoned that we might have more power to detect genetic effects using the ratio of *w*AlbA:*w*AlbB density because the two *Wolbachia* strains within an individual have a shared environment. However, again there was no significant effect (Table 1). In contrast, there were significant effects of population on both *w*AlbA and *w*AlbB densities as well as the ratio of *w*AlbA:*w*AlbB density (Table 1, Figure 3), and this was supported even when excluding populations with single samples (Sri-Lanka, Fiji and Vanuatu) and male specimens from the analysis. The eight males in the data had significantly lower *w*AlbA densities but no difference in *w*AlbB densities (Table 1, Figure 3). This is consistent with *w*AlbA having a high density in ovaries and *w*AlbB a wider tissue tropism (Ant and Sinkins 2018; Chuchuy et al. 2025).

**Table 1.** Linear models on *Wolbachia* density variation.

| Predictors | wAlbA density | wAlbB density | wAlbB/wAlbA ratio |
| --- | --- | --- | --- |
| subclade | $F_{4,74} = 0.689$<br>$P = 0.6$ | $F_{4,74} = 2.193$<br>$P = 0.08$ | $F_{4,74} = 1.236$<br>$P = 0.30$ |
| population | $F_{6,74} = 4.287$<br>$P = 0.0009$ | $F_{6,74} = 3.410$<br>$P = 0.005$ | $F_{6,74} = 5.274$<br>$P = 0.0001$ |
| sex | $F_{2,74} = 4.196$<br>$P = 0.02$ | $F_{2,74} = 0.720$<br>$P = 0.49$ | $F_{2,74} = 10.756$<br>$P = 0.0001$ |
| developmental stage | $F_{1,74} = 2.027$<br>$P = 0.16$ | $F_{1,74} = 3.604$<br>$P = 0.06$ | $F_{1,74} = 0.520$<br>$P = 0.47$ |

To quantify the roles of genes and the environment, we investigated whether related samples tend to have similar *Wolbachia* densities using a 2-strain bivariate phylogenetic mixed model. This allowed us to estimate the proportion of the variance in *Wolbachia* density that is explained by the phylogeny—the phylogenetic heritability. Regardless of whether ‘population’ is included in the model, the phylogeny explained a negligible proportion of the variance in density, with lower bounds close to zero (0.2-0.4 %, Table 2). Similar results were obtained when analysing the ratio of *w*AlbA:*w*AlbB densities using a univariate phylogenetic mixed model (Table 2). Nonetheless, while we do not find evidence for the *Wolbachia* genome affecting density, our credible intervals are large, so we equally do not find compelling evidence for a lack of genetic effects (Table 2). In contrast, population explained 32-35% of the variance in *w*AlbA density and 51-57% of the variance in the ratio of *w*AlbA:*w*AlbB densities but a negligible proportion of the variance in *w*AlbB. In the bivariate models, the *w*AlbA and *w*AlbB residuals were strongly correlated, consistent with unknown environmental factors affecting both strains Table 2. Therefore, non-genetic environmental effects are the most important determinants of the variation we observed in density.

**Table 2.** Phylogenetic mixed models on Wolbachia density. . Numbers in brackets indicate lower and upper bounds of credible intervals for estimated variance and correlation parameters.

| MCMCglmm models | Random effects | Two-strain models | wAlbB/wAlbA ratio models |
| --- | --- | --- | --- |
| phylogeny only | Phylogenetic heritability ($h^2$) | wAlbA: 0.004 [0.0001 ; 0.63]<br>wAlbB: 0.003 [0.0002 ; 0.38] | 0.003 [0.00009 ; 0.69] |
|  | wAlbA-wAlbB correlation of residuals | 0.57 [0.38 ; 0.71] | - |
| population only | Population | wAlbA: 0.35 [0.08 ; 0.76]<br>wAlbB: 0.003 [0.0001 ; 0.31] | 0.57 [0.15 ; 0.79] |
|  | wAlbA-wAlbB correlation of residuals | 0.67 [0.47 ; 0.74] | - |
| phylogeny + population | Phylogenetic heritability ($h^2$) | wAlbA: 0.002 [0.00004 ; 0.38]<br>wAlbB: 0.003 [0.0001 ; 0.42] | 0.001 [0.00001 ; 0.15] |
|  | Population | wAlbA: 0.32 [0.05 ; 0.72]<br>wAlbB: 0.003 [0.0001 ; 0.32] | 0.51 [0.13 ; 0.77] |
|  | wAlbA-wAlbB correlation of residuals | 0.61 [0.47 ; 0.75] | - |

### Plasmids are vertically transmitted within *w*AlbA

Two plasmids, pWALBA1 and pWALBA2, have been reported from *w*AlbA. To test whether plasmid abundance was more closely associated with *w*AlbA or *w*AlbB in our dataset, we fitted SMA regressions between each plasmid sequencing depth and each *Wolbachia* strain after removing variation explained by differences in mitochondrial depth (see methods, Figure 2C-D). Since *w*AlbA and *w*AlbB densities in the same individuals are correlated, we also performed partial SMA analyses in which the effect of one *Wolbachia* strain was removed from both the plasmid and the other *Wolbachia* strain before fitting the final regression. Using this approach, the sequencing depths of both plasmids were strongly associated with *w*AlbA, whereas the reciprocal associations with *w*AlbB were non-significant. This suggests that the plasmid copy number is tightly regulated and both plasmids have a strict association with *w*AlbA, as opposed to moving between the two strains. Furthermore, in most samples, pWALBA1 had a higher depth of coverage than pWALBA2 (x̄ = 1.64×, Figure S2), indicating that it usually has a higher copy number within *Wolbachia* cells. These results support earlier observations, where the plasmids were identified in *w*AlbA with ∼5-10 plasmids per *Wolbachia* genome, and with pWALBA1 generally having a higher copy number than pWALBA2 (Martinez, Ant, et al. 2022).

### *Wolbachia* and mitochondrial phylogenies are concordant

Horizontal or paternal transmission of *Wolbachia,* plasmids or mitochondria will result in the genomes having different tree topologies. Accordingly, we checked whether there were clades that were well-supported in one tree that conflicted with well-supported clades in another tree. The mtDNA, *w*AlbA, and *w*AlbB trees were all concordant across major clades (Figure 4; Figure S3). There were a small number of cases where clades that were well supported in one tree conflicted with well supported clades in another tree (e.g. Japan009, 013, and 015). However, we concluded these did not provide robust evidence of horizontal transmission. The conflicts were all between closely related strains, and inspection of the alignment revealed the conflicts were supported by very few sites within the genome. Furthermore, in some cases the derived SNP alleles causing these conflicts also occurred independently in distantly related strains or were very nearby in the genome, as might be expected if there was a bioinformatic error or high mutation rate at that position. The pWALBA1 and pWALBA2 genomes each had few variable sites but branches were consistent with vertical transmission of these plasmids (Figure S4).

**Figure 4.**
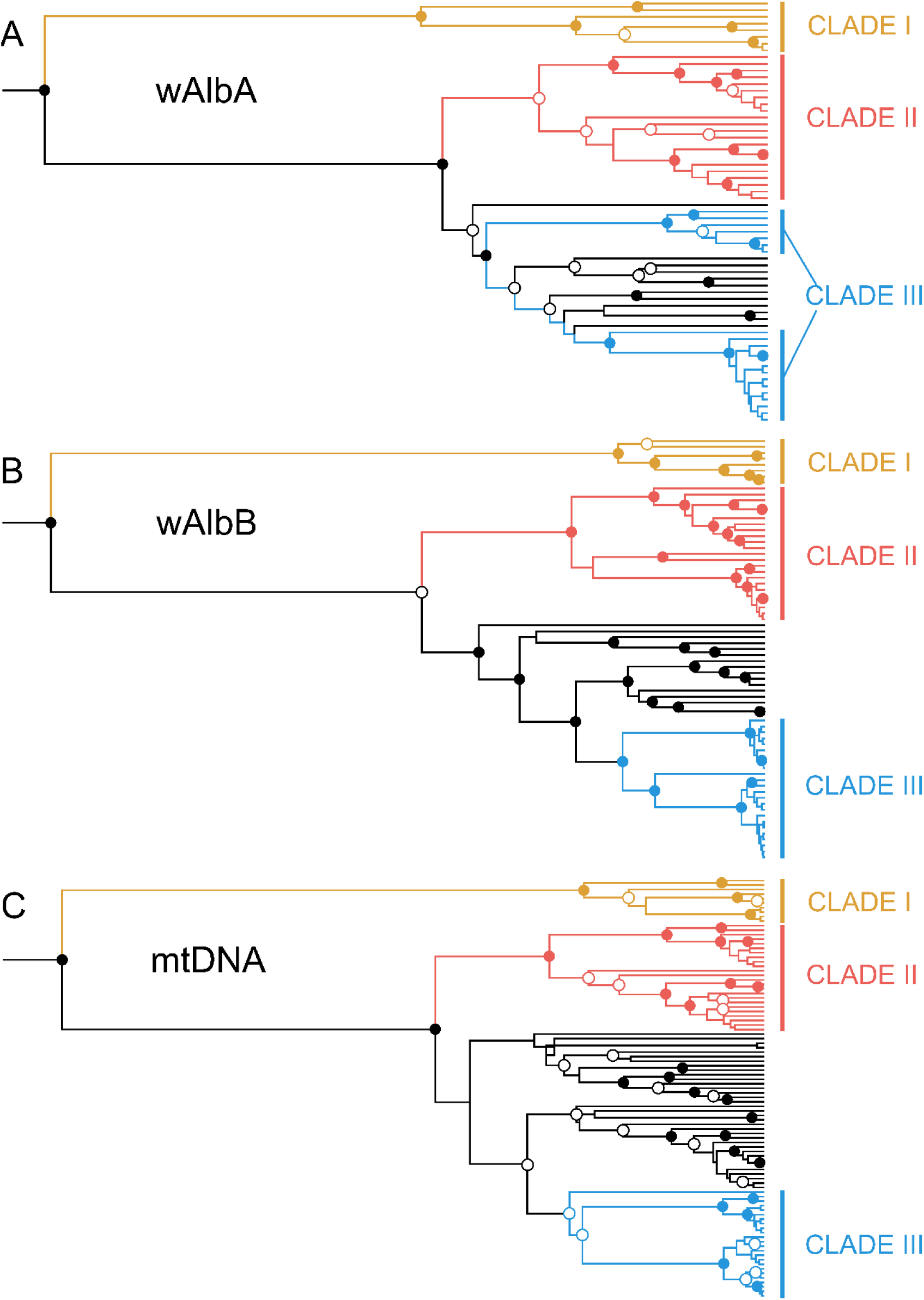
Phylogeny of the *Wolbachia* and mtDNA genomes. Maximum clade consensus trees constructed in Beast v1.10.4 using an HKY substitution model with a gamma site heterogeneity model, an uncorrelated relaxed clock and a constant size coalescent tree prior (Suchard et al. 2018). Independent substitution models were set for 1^st^ codon positions, 2^nd^ codon positions, 3^rd^ codon positions, non-coding RNA, and non-coding intergenic DNA. Filled circles indicate nodes with ≥0.95 posterior support, white circles indicate 0.5 – 0.95 support. Three major clades described in the main text are indicated by color. Figure S3 is an identical plot with sample names and support values.

### Evolutionary relationships of *Wolbachia* strains reflect the history of mosquito populations

There was strong spatial genetic structure within the data, with some populations forming monophyletic clades on the *Wolbachia* and mtDNA trees (Figure S3). As trees of the different genomes were largely concordant, we also built a composite phylogeny using the 59 samples for which we had all five genomes. The resultant tree had strong support for all but the most terminal clades and also showed strong geographical structure (Figure 5A). All the trees had a single deep split, separating a group containing Timor-Leste and half of the Torres Strait Islands (TSI) samples (Clade I) from all other samples (Figures 4,5A,S3). As the TSI was only invaded by *Ae. albopictus* in 2004 (Ritchie et al. 2006), and has a nuclear genomic background predominantly derived from Indonesia/Timor (Beebe et al. 2013; Maynard et al. 2017; Schmidt et al. 2021), Clade I is likely a lineage found across Indonesia and Timor. Notably, the differentiation of this lineage was greater than between putative native range populations in mainland Southeast Asia (Vietnam, Malaysia) and populations in East Asia that were likely invaded far before the recent 20^th^ century range expansion (China, Japan). Within the TSI was a second lineage related to lineages in Papua New Guinea (PNG) and Malaysia (Clade III) that was intermingled with and highly divergent from the Indonesian/Timorese lineage (Clade I) (Figures 5A,S3). This points to an additional genetic source of the TSI invasion, from either PNG or a genetically similar region, that either admixed with Clade I in southern PNG prior to invasion or spread into the TSI via gene flow following invasion; either hypothesis accords with nuclear genomic patterns (Schmidt et al. 2021; Schmidt et al. 2024).

**Figure 5.**
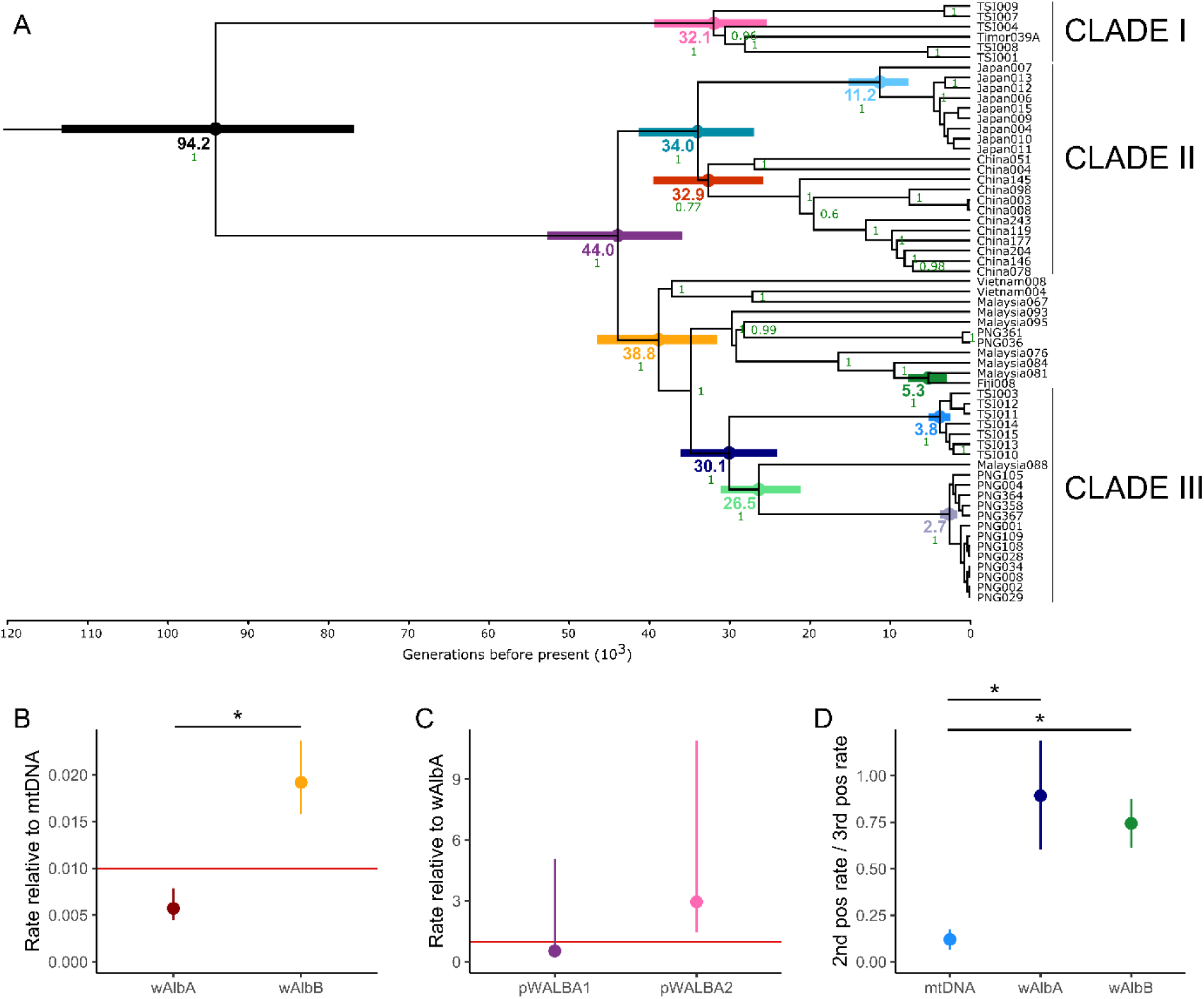
Phylogeographic dating of node ages and calibration of evolutionary rates. (A) Combined tree for the 59 strains with all five genomes constructed in Beast v1.10.4, using an HKY substitution model with a gamma site heterogeneity model, a strict molecular clock and a constant size coalescent tree prior (Suchard et al. 2018). Independent substitution models were set for 1^st^ codon positions, 2^nd^ codon positions, 3^rd^ codon positions, non-coding RNA, and non-coding intergenic DNA. Numbers at nodes indicate posterior support, plotting only values ≥0.5 and with some values at tips not plotted for clarity. Three major clades described in the main text are indicated on right. Horizontal boxplots indicate the 95% highest probability density (HPD) range for node ages. (B) HPDs and means of short-term evolutionary rates of *w*AlbA and *w*AlbB 3^rd^ codon positions relative to mtDNA. Red horizontal line indicates the rate of *w*Mel relative to *D. melanogaster* (Richardson et al. 2012). (C) HPDs and means of short-term evolutionary rates of pWALBA1 and pWALBA2 3^rd^ codon positions relative to *w*AlbA. Red horizontal line indicates equal rates. (D) HPDs and means of short-term evolutionary rates at 2^nd^ codon positions relative to 3^rd^ codon positions. Asterisks (*) indicate where 95% HPD intervals of differences between posterior distributions do not overlap zero. TSI = Torres Strait Islands; PNG = Papua New Guinea.

A second early split in the trees separates the East Asian samples of China and Japan (Clade II) from the remaining samples (Figures 5A,S3). This latter group was roughly split into samples from PNG and the TSI (Clade III) and those from Vietnam, Malaysia, and Fiji. PNG, Vietnam and Malaysia were sometimes intermingled with other populations, reflecting gene flow or ancestral polymorphisms. Within Malaysia, samples from Pahang (east) and Johor (south) did not cluster by location.

The publicly available genomes of *w*AlbB-Uju and *w*AlbB-Hou are well-characterised, and these strains have diverged both in gene content and traits likely to affect the success of release programs (Martinez, Ross, et al. (2022)). The Italian *w*AlbB-Uju strain was in Clade II (Figure S3B). The American *w*AlbB-Hou strain lacked a clear placement for *w*AlbB (Figure S3B), but its mtDNA genome was most related to a Vietnamese strain that was filtered from the wAlbB analysis (Figure S3C). A third publicly available sequence from Malaysia (*w*AlbA-JF2017) clustered with other Malaysian samples (Figure S3C). Notably, only *w*AlbB-Uju and *w*AlbB-Hou have been investigated for *Ae. aegypti* release programs, but there is much unexplored genetic variation. These two variants differed by 191 SNPs (c.f. 130—141 SNPs in Martinez, Ross, et al. (2022)), while they were each almost twice as differentiated from Clade I (338 and 378 SNPs respectively).

### The *Wolbachia* superinfection has a recent common ancestor in Southeast Asia

To estimate divergence dates, we used the composite phylogeny of the 59 samples for which we had all five genomes (Figure 5A). In *Drosophila* populations, we have previously calibrated the rate of molecular evolution using the mtDNA mutation rate derived for *D. melanogaster* (Haag-Liautard et al. 2008; Richardson et al. 2012). We used this as a prior for mtDNA 3^rd^ codon substitution rates in *Ae. albopictus* to estimate divergence times and evolutionary rates in this study, setting a strict molecular clock (Table S2). Following Richardson et al. (2012), these rate estimates are neither mutation rates (which require observation of strongly deleterious mutations) nor long-term substitution rates (which would exclude slightly deleterious mutations), but instead represent short-term microevolutionary rates.

We estimate that all samples share a common ancestor 76,827—113,520 generations before present (x̄ = 94,233; Figure 5A). This represents the earliest split in the dataset, between the Indonesian/Timorese lineage of Clade I (Figure 4) and all other samples. The East Asian populations in Clade II split from Southeast Asian populations 35,801—52,899 (x̄ = 43,954) generations ago, followed by more recent splits among Malaysia and Vietnam and the presumed invasive range populations. Converting generational estimates to years is non-trivial in *Aedes* mosquitoes given that generation times vary according to local climate, where locations with pronounced dry seasons (e.g. TSI) should have fewer gens/year than those without (e.g. Malaysia). Shorter generation time might also be selected for during the range expansions following invasion, which may disproportionately affect estimates from recently invaded populations. Nevertheless, *Aedes* generation times are commonly assumed to range from 10 to 15 gens/year (Crawford et al. 2017; Rose et al. 2023). Assuming 10 gens/year dates the common ancestor of all samples at 7,683—11,352 years before present (x̄ = 9,423), while assuming 15 gens/year dates it at (5,122—7,568) years before present (x̄ = 6,282).

### *Wolbachia* genomes evolve at different rates

The two *Wolbachia* strains had markedly different evolutionary rates from mtDNA and from each other (Figure 5B; Table S2). Across all codon positions, *w*AlbA evolved slower than *w*AlbB, suggesting that it has a lower mutation rate. Both genomes evolved far slower than mtDNA, where the 3^rd^ codon rate was 127—224× faster (x̄ = 175×) than *w*AlbA and 42.2—63.3× faster (x̄ = 52.1×) than *w*AlbB. These estimates fall either side of the equivalent statistic in *D. melanogaster*, where the 3^rd^ codon rate was ∼100× faster in mtDNA compared to the *w*Mel *Wolbachia*. These results appear to justify our application of the *D. melanogaster* rate prior to *Ae. albopictus*, as the evolutionary rates of mtDNA and *Wolbachia* have similar ratios across both systems. The *w*AlbA 3^rd^ codon rate was 0.197—4.98× (x̄ = 1.86×) the rate of pWALBA1 but only 0.0916—0.681× (x̄ = 0.338×) the rate of pWALBA2 (Figure 5C), though these comparisons were strongly limited by the number of variable sites for inference.

The evolution of the mitochondrial genome was dominated by purifying selection, with third codon positions evolving at over eight times the rate of second codon position (changes at second codon positions normally alter the amino acid, while those at third codon positions do not) (Figure 5D). In contrast, in both *Wolbachia* genomes there was only a slight difference in rates between the two classes of sites (Table S2). This indicates purifying selection is a weak force on these *Wolbachia* genomes.

### Widespread variation in *w*AlbB gene content

We estimated the copy number of each gene from the depth of sequencing coverage, then focused on genes that had copy number = 0 (i.e. missing) in at least 5 samples or showed copy numbers >2 in at least one sample. Applying these threshold values to *w*AlbB samples, we found a total of 184 genes that varied in presence/absence or copy number, including genes located in prophage regions and accessory genes of the phage eukaryotic association module (Figure 6A, Table S3). While the gene content profiles mostly reflected the *w*AlbB phylogeny, some clusters of genes were lost independently in distantly-related clades, such as the prophage region encoding proteins of the phage tail fibre (DEJ70_05885-DEJ70_05900). In the *w*AlbB-Hou reference genome, this prophage region is flanked by nearly identical insertion sequences encoding IS481 family transposases which likely facilitated its loss in multiple *w*AlbB clades. In the fully assembled *w*AlbB-Hou and *w*AlbB-Uju genomes, the WO prophage has rearranged into several regions and lost multiple essential genes, suggesting that the phage genome is too degraded to undergo phage replication. However, the exact genomic architecture of prophage regions in our *w*AlbB samples is uncertain because of regions absent from the *w*AlbB-Hou genome that was used to design our capture probes were not sequenced.

**Figure 6.**
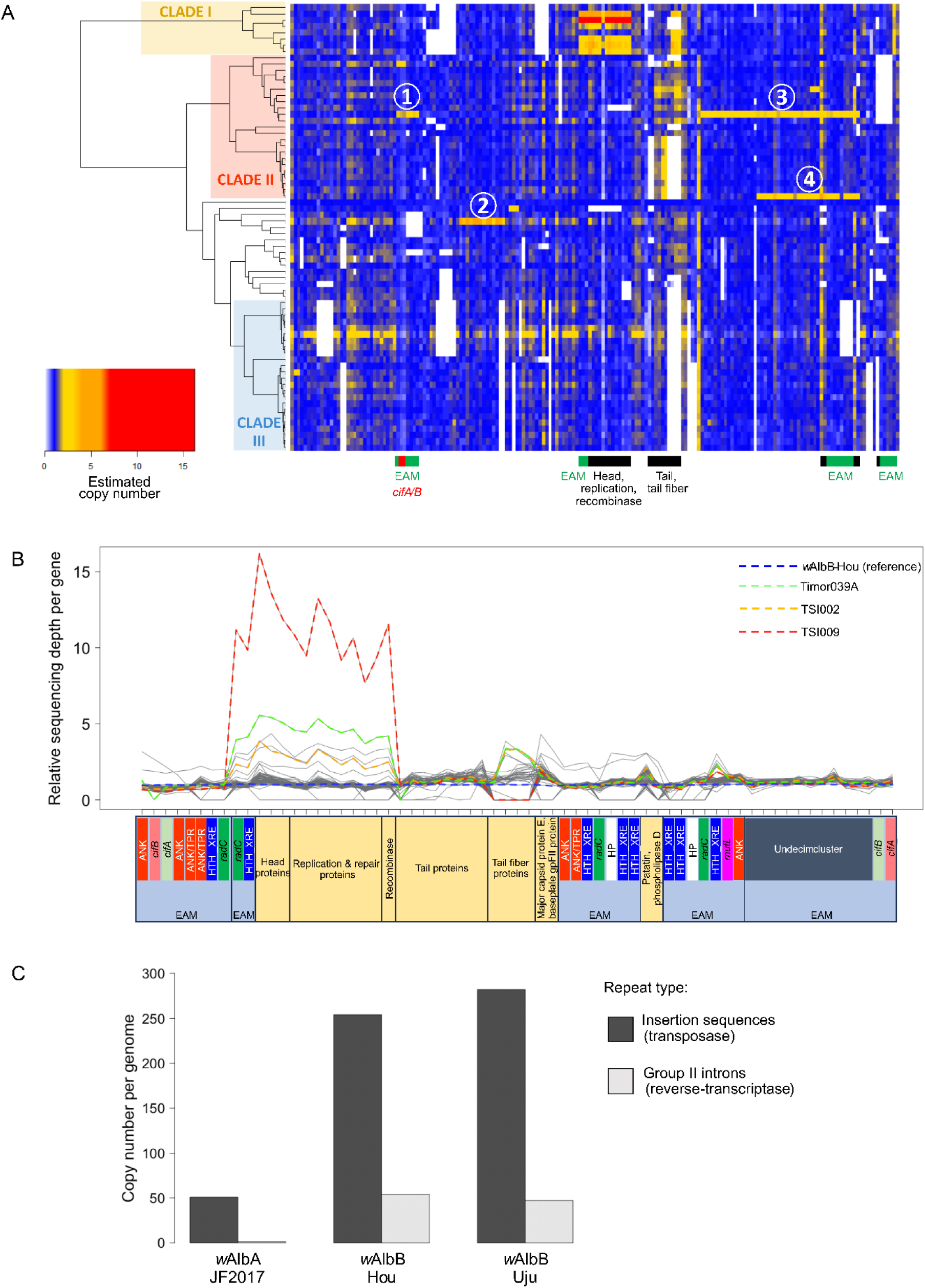
Gene content variation. (A) Left: *w*AlbB phylogeny. Right: Heatmap of estimated copy number per gene (columns) and per sample (rows), Estimated copy number was the median sequence depth of a given gene divided by the mean across all genes in the genome (excluding genes mapping to multiple genomic locations). White circles with numbers (1–4) indicate putative large duplications: 1 (24.7 kb), 2 (25 kb), 3 (127 kb) and 4 (92 kb). Colored bars at the bottom indicate genes located in prophage regions: black = phage essential genes; green = accessory genes of the eukaryotic association module or EAM; red = cytoplasmic incompatibility factors or *cif* genes. (B) Estimated gene copy numbers in *w*AlbB prophage regions. Dashed colored lines indicate three samples of interest and the reference genome. ANK/TPR: ankyrin/tetratricopeptide repeat-containing gene; *cifA/B*, cytoplasmic incompatibility factors A and B; HTH_XRE, helix-turn-helix transcription regulator; *radC*, RadC-like DNA repair protein; *mutL*, DNA mismatch repair protein MutL; Undecim cluster, putative “pathogenicity island” involved in host-symbiont interactions; HP, hypothetical protein. (C) Repeat content of *w*AlbA and *w*AlbB references genomes. IS elements were predicted using ISEScan (Xie and Tang 2017) and group II introns counted from CP101657.1, CP031221.1 and CP102671.1 annotation files available on Genbank.

*w*AlbB genomes in samples from the divergent Clade I showed an unusually high copy number of a prophage region encoding recombinase, head proteins and proteins involved in phage replication (DEJ70_04880-DEJ70_04945, Figure 6A-B and Table S3). While this could indicate that the region is amplified within the *Wolbachia* genome, the >5–15× higher sequencing depths in some samples suggests that this is unlikely. Instead, such high copy numbers may be due to the partial replication of a phage genome, with other phage genes not being replicated perhaps due to genomic rearrangements of the complete prophage region (Figure 6B). Other prophage modules such as the tail fibre region and parts of the eukaryotic association module displayed ∼2–3× increases in sequencing depth in some samples which may be indicative of genomic duplications (Figure 6B).

We observed four other putative large duplications in samples China078, Malaysia059 and Japan013 (numbered circles in Figure 6A) showing ∼2× to 4× amplification (Figure S5A-C). In the *w*AlbB-Hou reference genome, these regions are ∼25 to 127 kb long and numerous repeat elements (transposase, reverse transcriptase) are found within them and/or in the flanking regions. This indicates that *w*AlbB genome size may vary substantially between samples.

Cytoplasmic incompatibility is induced by a pair of genes called *cifA* and *cifB*, with two divergent pairs found in the WO phage eukaryotic association module of *w*AlbB (Figure 6B). One of the *cifA-B* pairs has been independently duplicated in China078 (Duplication 1) and China145 (Figure S5A). Conversely, the cytoplasmic incompatibility toxin-encoding *cifB* gene was absent in Timor039A (Figure S5A). This gene is only required to induce cytoplasmic incompatibility in males, and theory predicts that because of this it can be lost without incurring any fitness cost, even when other strains in the population carry functional genes (Martinez et al. 2021). Six SNPs were identified in the two pairs of *cifA-B* genes found in *w*AlbB-Hou genome, three of which were non-synonymous and located in the *cifB* copy DEJ70_07095 (Table S4). These included a Phe – Leu allele found in Vietnam and Malaysia, a Gly – Val allele found in Malaysia and Fiji, and an Ala – Val allele found in China.

Given that the *w*AlbB-Hou and *w*AlbB-Uju variants are known to differ in symbiont density depending on temperature and host background (Martinez, Ross, et al. 2022), we tested for correlations between gene copy number and *w*AlbB density within samples. Among the variable genes, none of them were significantly associated with *Wolbachia* density after correcting for multiple testing (Kruskal-Wallis test per gene with three states: “absent” versus “1 copy” versus “> 1 copy”; Table S5).

### *w*AlbA gene content and plasmids

In contrast with *w*AlbB, our sequencing depth analysis did not reveal any large deletions or duplications among *w*AlbA samples. One explanation for this striking difference in genome plasticity between the two strains could be that they differ in their repeat content. We investigated repeat content using the *w*AlbA and *w*AlbB reference genomes, as these were assembled from long reads as well as short reads. We found evidence of greater repeat content in *w*AlbB (Figure 6C). Indeed, both *w*AlbB-Hou and *w*AlbB-Uju circular assemblies contain much more insertion sequences and group II intron reverse-transcriptases than *w*AlbA-JF2017, suggesting that these repeat elements could be promoting large chromosomal rearrangements across *w*AlbB lineages (Figure 6A).

Despite the lack of large genomic changes between *w*AlbA samples, there were 68 genes with copy numbers >1 in at least one sample, though these did not follow any clear genomic distribution or phylogenetic structure (Figure S6A). *w*AlbA sequencing depths were substantially noisier than *w*AlbB across the genome, (Figure S6B-C), and we considered the signal-to-noise ratio to be too low to estimate *w*AlbA gene copy number with accuracy. This may in part be because the *w*AlbA DNA-capture probes were designed on an incomplete and highly fragmented genome assembly, which was not the case for *w*AlbB. We observed little evidence of variation in gene copy number in the two plasmids pWALBA1 and pWALBA2, with none of the plasmid genes showing relative copy number >2 and no genes absent in any sample (Figure S7; two copies of a transposase in pWALBA2 in the reference genome were not estimated due to multiple mapping of the reads).

## Discussion

*Wolbachia* superinfections are common in nature (Sinkins et al. 1995; Riegler and Stauffer 2002; Watanabe et al. 2011) and transinfected superinfections are being used in mosquito control programs (Zheng et al. 2019). The *w*AlbA*–w*AlbB superinfection of *Aedes albopictus* provides an opportunity to examine how long-term vertical transmission shapes the evolution of co-inherited symbionts in a major disease vector. Our data show that this superinfection has persisted for approximately 6,000–9,000 years with no evidence of horizontal transfer, and that its global distribution has largely tracked the demographic history of its host. However, shared transmission has not produced uniformity: *wAlbA* and *wAlbB* differ in evolutionary rate, and *wAlbB* shows extensive structural variation. Infection density varies over orders of magnitude among individuals and appears to be driven predominantly by environmental rather than phylogenetic effects. Therefore, vertical inheritance can maintain long-term genomic associations among host and symbionts, and this has been accompanied by substantial diversification in symbiont genome content and infection phenotype.

Our estimate that the current *w*AlbA-*w*AlbB superinfection likely originated within the past ∼6000-9000 years (approximately 77,000—114,000 mosquito generations ago) aligns with a previous, less precise estimate from mitochondrial COI sequences (1,790—27,600 ybp; x̄=9,040 (Maynard et al. 2017). This is similar to *w*Mel strains in *D. melanogaster*, which were estimated to have a common ancestor ∼8,000 years ago (Richardson et al. 2012). Our divergence dates may mark the onset of superinfection in *Ae. albopictus* following the introduction of one or both *Wolbachia* strains from another host species. It has long been known that *Wolbachia* infections are often transient within host populations and are sustained through horizontal transfers between species, and recent origins of infections of the *D. melanogaster* and *Ae. albopictus infections* aligns with analyses of different species of *Drosophila* which identified multiple horizontal transfer events within the last 14,000 years (Turelli et al. 2018; Turelli et al. 2022). Alternatively, the dates could represent the fixation of a particular lineage within an older infection, driven by a selective sweep or genetic drift.

Comparing the *Wolbachia* and mitochondrial phylogenies, we found no evidence of horizontal or paternal transmission of *w*AlbA or *w*AlbB. These modes of transmission would break down the association between *Wolbachia* and mtDNA in a manner akin to recombination breaking down linkage disequilibrium between genes on the same chromosome. Even low rates of recombination lead to the rapid decay of linkage disequilibrium, so if any horizontal or paternal transmission of *Wolbachia* occurring, the rate is likely to be extremely low. This aligns with population genetic and experimental work in other species (Charlat et al. 2009; Richardson et al. 2012; Bakovic et al. 2020), and confirms that, while horizontal transmission is key in host jumps and long term evolution, it is unlikely to have any effect on the spread of *Wolbachia* within populations of these species.

Two lines of evidence suggest that the superinfection has reached its present distribution by co-migrating with *Ae. albopictus* as it invades new regions, rather than spreading through existing mosquito populations. The first is that the nuclear genetic structure in *Ae. albopictus* is strikingly similar to the phylogenetic structure of the *w*AlbA—*w*AlbB superinfection (Schmidt et al. 2024). This aligns most plausibly with the co-migration hypothesis. The second is that, while the *w*AlbA—*w*AlbB superinfection induces cytoplasmic incompatibility in crosses to singly infected mosquitoes (Dobson et al. 2004), which can maintain the infection at high frequencies within populations, laboratory experiments failed to detect any direct fitness advantage of superinfections over single infections of either strain (Dobson et al. 2004). Without a fitness advantage to the superinfection, it would be difficult for superinfection to spread across the Asian range of *Ae. albopictus* in the 6000-9000 years since its origination, as any imperfect transmission or local fitness costs result in a threshold frequency that *Wolbachia* must exceed to invade a population (Hoffmann et al. 1990). Work in *Aedes aegypti* has shown that even modest barriers to mosquito dispersal can prevent the spatial spread of *Wolbachia* under these conditions (Schmidt et al. 2017). Together, these observations suggest that the superinfection has most likely reached its present distribution by co-migrating with *Ae. albopictus* as it invades new regions.

If this hypothesis is correct, it points to a native range of *Ae. albopictus* in Southeast Asia but not East Asia. The deep phylogenetic split between Indonesia/Timor-Leste (Clade I) and other regions suggests a specific origin in Indonesia/Timor-Leste or in nearby Mainland Southeast Asia. It also suggests that the current and ongoing global expansion of *Ae. albopictus* beginning in the 19^th^ and 20^th^ centuries was predated by a much earlier expansion into East Asia ∼2,500—5,000 years ago. This finding aligns with some earlier predictions (Paupy et al. 2009) but clashes with studies that have assumed an origin in East Asia (Bonizzoni et al. 2013; Sherpa et al. 2022).

Although the superinfection was present in every population, *Wolbachia* density varied by >4 orders of magnitude across individual mosquitoes. This is significant as *Wolbachia* density is a major driver of its antiviral effects, and influences transmission and other phenotypes (Chrostek et al. 2013; Martinez et al. 2014). To verify that samples with low *Wolbachia* density represented genuine infections, we examined raw sequencing reads and confirmed that the genetic variants in the *Wolbachia* genomes mostly matched those expected based on the mitochondrial background, supporting the conclusion that these low-density infections are real. However, we caution that these infections may be functionally unimportant. First, we are detecting *Wolbachia* DNA but have no evidence that the bacteria are viable. Second, the densities may be so low that *Wolbachia* is neither transmitted nor capable of affecting host phenotype such as through cytoplasmic incompatibility. This may depend on the tissue tropism of these low-density infections; if *Wolbachia* is concentrated in the ovaries, even at low overall densities, it may still be transmissible or induce cytoplasmic incompatibility (Schneider et al. 2018; Hague et al. 2024). Overall, the questions raised by our study around the biological importance of low-density infections are critically important given these same samples were often scored as uninfected using qPCR (Yang et al. 2022). If low-density infections are transmissible and/or induce phenotypic effects on hosts, population-level screening for *Wolbachia* with qPCR will lead to systematic underestimation of infection frequencies and inaccurate inference of associated parameters including those central to *Wolbachia*-based pest management strategies (Schmidt et al. 2017; Turelli and Barton 2017).

Our analysis demonstrated that environmental factors are the primary drivers of variation in *Wolbachia* density. While much of the variation is seen within populations, there are also substantial differences between different geographical locations. The observation that local environment controls density may present challenges for the effectiveness of control programs in certain regions. However, it should also be kept in mind that relative intracellular density and associated phenotypes of *w*AlbB variants following transfer into *Ae. aegypti* may be different to those observed in their native host *Ae. albopictus*.

We failed to find evidence of genetic effects when we tested whether related *Wolbachia* strains tend to exhibit similar symbiont densities. However, our estimates were very uncertain, so we may have failed to detect genetic effects due to the large environmental variation. Furthermore, genetic effects may only become apparent when taking into account genotype-environment interactions (Hague et al. 2022; Martinez, Ross, et al. 2022). Our previous work showed that *w*AlbB-Hou and *w*AlbB-Uju show similar densities under standard conditions but differ significantly at very high temperatures, the magnitude of this effect being also dependent on the mosquito genetic background (Martinez, Ross, et al. 2022). Therefore, it will be necessary to experimentally disentangle the contributions of genetic and environmental factors to the variation in density, by comparing lab colonies of different *w*AlbB variants in a common host genetic background and under controlled conditions. If *Wolbachia* genetic factors are significant, our findings could inform the selection of optimal *Wolbachia* strains for use in vector control programs.

Despite their recent common ancestry, we observed a surprising degree of structural variation between *w*AlbB variants, including changes that affect gene content. As seen in other *Wolbachia* strains (Cerveau et al. 2011; Leclercq et al. 2011; Amoros et al. 2026), this variation appears to be frequently driven by mobile genetic elements such as prophages and short insertion sequences. In some cases, apparent differences in gene copy number may reflect ongoing phage replication rather than genomic rearrangements. The persistence of this variation within populations may be facilitated by weak or ineffective purifying selection (see below). A notable example is the *cifB* gene, which is responsible for modifying sperm to induce cytoplasmic incompatibility (LePage et al. 2017). Since *Wolbachia* is maternally inherited, evolutionary theory predicts that selection to maintain *cifB* weakens once the symbiont becomes fixed in a population (Turelli 1994; Martinez et al. 2021). Consistent with this prediction, we detected a deletion of *cifB* at low frequency. If this spreads through populations, it may be the first step towards the loss of the infection.

To date, only two *w*AlbB strains—*w*AlbB-Hou and *w*AlbB-Uju—have been introduced into *Ae. aegypti* for dengue control. These strains originate from recently invaded regions (the USA and Italy, respectively) and exhibit clear phenotypic differences (Martinez, Ross, et al. 2022). In this study, we identify a divergent lineage (Clade I) with an Indonesian origin that is nearly twice as genetically distant from the two released strains as they are from each other. Clade I also had differences in copy number and gene loss from other lineages. Among this global diversity of *w*AlbB, there is likely to be a broad range of phenotypes with relevance to dengue control. This raises the possibility that currently deployed strains may be suboptimal compared to unexplored alternatives (Salje and Jiggins 2024).

The shared evolutionary history of *Wolbachia*, plasmid, and mitochondrial genomes enables direct comparisons of their evolutionary rates. Third codon positions, where mutations are mostly synonymous and thus relatively unconstrained, evolve ∼100× more slowly in *Wolbachia* than in mitochondrial DNA (mtDNA), indicating a substantially lower mutation rate in *Wolbachia*. A similar disparity was observed between *w*Mel and *D. melanogaster*, supporting the hypothesis that *Wolbachia* possesses efficient DNA repair mechanisms (Richardson et al. 2012). Accordingly, the observation that *w*AlbB evolves ∼3.4× faster than *w*AlbA at third codon positions may reflect the higher densities of *w*AlbB within superinfected hosts, leading to more frequent replication and a higher mutation rate. Interestingly, the evolutionary rate at second codon positions—where mutations are largely nonsynonymous—was only ∼25% lower than at third positions. This suggests that *Wolbachia* protein-coding sequences are evolving with relatively little selective constraint. A similar pattern in *w*Mel implies this may be a general feature of *Wolbachia* genomes (Richardson et al. 2012). In contrast, second codon positions in mtDNA evolve much more slowly than 3^rd^ codon positions, reflecting stronger purifying selection. Given that these genomes are in near-perfect linkage disequilibrium, the elevated rate of protein evolution in *Wolbachia* may result from relaxed selective constraints, at least relative to the mitochondria. However, genome-wide comparisons of *Wolbachia* strains reveal non-synonymous substitution rates are approximately one fifth of the synonymous substitution rate, suggesting that over longer evolutionary timescales, many nonsynonymous mutations are purged, and the observed high rates may reflect recent mutations not yet filtered by purifying selection (Rocha et al. 2006; Richardson et al. 2012).

## Conclusion

Our analyses show that the *w*AlbA*–w*AlbB superinfection of *Aedes albopictus* has persisted through strict maternal transmission for ∼6,000–9,000 years, with its spread tightly coupled to host population history. Over this timescale, the two symbionts’ genomes have evolved in markedly different ways: *wAlbA* shows slower rates of nucleotide sequence evolution, whereas *wAlbB* has undergone extensive structural change, including recurrent variation in prophage regions and cytoplasmic incompatibility genes. Symbiont density varies over several orders of magnitude among individuals and is only weakly associated with symbiont ancestry, indicating a predominant role for environmental effects. These results show that long-term vertical transmission does not imply genomic or phenotypic stability in *Wolbachia*. Instead, natural populations harbour substantial diversity at levels likely to affect both symbiont biology and the performance of *Wolbachia*-based vector control.

## Materials and Methods

### Sample selection

The 94 samples previously screened for *Wolbachia* infection and sequenced with ddRADseq were extracted using either Qiagen DNeasy Blood & Tissue Kits (Qiagen, Hilden, Germany) or Roche High Pure™ PCR Template Preparation Kits (Roche Molecular Systems, Inc., Pleasanton, CA, USA). Additional, previously-sequenced samples were from Kuala Lumpur (*w*AlbA-JF2017; Martinez, Ant, et al. (2022)), the USA (*w*AlbB-Hou; Sinha et al. (2019)), and Italy (*w*AlbB-Uju; Martinez, Ross, et al. (2022)). The Italian strain (*w*AlbB-Uju) is named after an Indonesian location but does not have an Indonesian *w*AlbB background as the Indonesian-sourced mosquitoes were later backcrossed into an *w*AlbB-infected Italian background (Blagrove et al. 2012) and the *w*AlbB infection was transferred into *Ae. aegypti* before its genome was sequenced (Ant et al. 2018).

### Library preparation and target enrichment

DNA sequencing libraries were prepared using the NEBNext® Ultra™ II FS DNA Library Prep Kit for Illumina (New England Biolabs, E7805L) and NEBNext® Multiplex Oligos for Illumina® (New England Biolabs, E7335G) according to the manufacturers’ recommendations with variations as follows: all samples were fragmented for 10 minutes at 37°C. A post-ligation size selection step was performed only on libraries made from greater than 100ng genomic DNA starting material. For the size selections 25µl of NEBNext Sample Purification beads were used for the first step and 10ul of beads for the second step. The number of cycles of PCR enrichment of adaptor-ligated libraries was optimised to yield greater than 1µg of DNA in a pool for subsequent target captures. Between 6 and 12 PCR cycles were used, depending on the concentration of the initial genomic DNA starting material. The libraries were quantified using a Qubit HS DNA assay kit (Thermofisher Q32851). The relative concentration of each library was assessed using qPCR: 3 replicates of each library was diluted 1000-fold in 1ml of 10mM Tris pH 8.0, 0.05% Tween20. For each replicate a qPCR reaction was set up containing 5ul of 2x Sensifast Hi-Rox Sybr (Meridian BIO-92020), 2µl of nuclease free water, 1µl of 2.5µM of each library qPCR primer and 2µl of diluted library sample in a 384-well MicroAmp Optical Reaction Plate (Applied Biosystems 4309849). qPCR primers were: IS5_reamp.P5 AATGATACGGCGACCACCGA; and IS6_reamp.P7 CAAGCAGAAGACGGCATACGA. Plates were sealed using MicroAmp Optical Adhesive film (Applied Biosystems 4311971) vortexed to mix and centrifuged briefly. The plates were run on a 384-well QuantStudio™ 5 Real-Time PCR System (Applied Biosystems A28575) using the pre-set comparative-CT-Sybr with melt curve program with the reaction volume adjusted to 10µl. Data was analysed using QuantStudio^TM^ Design and Analysis Software v1.5.2. The CT threshold value was set to 0.1 for each plate.

Equimolar quantities of each library were then combined into 3 pools of 24 and 1 pool of 22. Pooled samples were enriched for either *Wolbachia* or mitochondria target DNA using a KAPA Hypercapture kit (Roche 09075810001) and KAPA Hypercapture Bead kit (Roche 09075780001) with custom KAPA Hyperexplore Max probes (Roche 09063633001) according to the manufacturer’s recommendations. A second probe for sequencing the pWALBA1 and pWALBA2 plasmids was designed along the same lines as the above and run with identical protocols. Probe design used the following published genomes as templates: *w*AlbA draft genome (GenBank accession NWVJ00000000); *w*AlbB-Hou (CP031221.1); mtDNA (KR068634.1); pWALBA1 (NZ_CP101658.1), pWALBA2 (NZ_CP101659.1). Where alternative options in the protocol were available, 20µl of COT human DNA (1mg/ml) was used to block the hybridisation reaction and 10 cycles of post-capture PCR amplification was used. Captures were eluted from the final bead wash in 20ul of EB, quantified using Qubit HS DNA assay kit and a 1:10 dilution was analysed using a Bioanalyzer HS DNA chip (Agilent, 5067-4626). Captures were quantified using qPCR as above and equimolar amounts were pooled together before sequencing using 2 lanes on the Illumina NovaSeq platform at the Cancer Research UK Cambridge Institute.

### Sequence processing, genotyping and filtering

Sequences were concurrently aligned to five genomes: *Ae. albopictus* mtDNA (Zhang et al. 2016), *Wolbachia* strains wAlbA (Martinez, Ant, et al. 2022) and *w*AlbB (Sinha et al. 2019), and plasmids pWALBA1 and pWALBA2 (Martinez, Ant, et al. 2022). Alignment was with bowtie2 (Langmead and Salzberg 2012) using –very-sensitive settings. Attempts to align with bwa mem (Li 2013) were unsuccessful due to frequent misalignment among the two *Wolbachia* strains. Following alignment, we used samtools v1.7 (Li et al. 2009) to generate sorted bam files, then Picard v2.27.4 to add read groups and mark duplicates, then samtools to retain only reads mapped as primary alignments in proper pairs.

Genotyping of samples was with GATK v4.2.6.1 (Auwera and O’Connor 2020). HaplotypeCaller was first used to produce gVCF files for every sample, setting EMIT_ALL_CONFIDENT_SITES. GenomicsDBImport was used to build these into a database. GenotypeGVCFs was used to genotype all samples concurrently, with settings to “--add-output-vcf-command-line”, “--include-non-variant-sites”, and “--sample-ploidy 1”. Hard filtering was applied to variants as follows: SelectVariants to remover indels; VariantFiltration for standard GATK hard filters (QD < 2.0, SOR > 3.0, FS > 60.0, MQ < 40.0, MQRankSum < -12.5, ReadPosRankSum < -8.0); and SelectVariants to “--set-filtered-gt-to-nocall TRUE”.

### *Wolbachia* density analysis

Since there was no clear separation in *Wolbachia* sequencing depth between infected and uninfected samples, we conducted a k-mer analysis of *w*AlbA and *w*AlbB SNPs across all samples in order to test if low depth of either *Wolbachia* strain was due to sequencing / bioinformatics artefacts, such as cross-sample contamination, or if it corresponds to genuine *Wolbachia* infections. To this end, we first selected a list of SNPs that were present in all samples showing a sequencing depth for either *Wolbachia* genome above 5x. This was done to minimize the presence of false-positive variant sites from low depth samples. From this set of reliable SNPs, we created a k-mer database by extracting from the two reference *Wolbachia* genomes respectively 10 and 9 bp on each side of every variable position, generating for each SNP one 20 bp k-mer carrying the reference allele and a corresponding k-mer with the alternate allele. We then counted in every sample how many times each k-mer in the database appears in the raw Illumina reads, using a custom python script. The ratio between reference and allele counts at each position and for each sample was used to calculate pairwise Euclidean distances between samples and determine the closest relative. Finally, we compared the mitochondrial lineage between target samples and their closest relative based on *Wolbachia* k-mers, with the prediction that they should match if *Wolbachia* infections in the two samples are real. The accuracy of this method was tested by randomly permuting (n = 10,000) the mitochondrial lineages between samples and comparing the distribution of matches in the permuted datasets to the observed number.

*Wolbachia* density was estimated by normalizing the genome depth of either *Wolbachia* strain to the depth of the mitochondrial genome in a given sample. Since *Wolbachia* and plasmid DNA libraries were prepared and sequenced separately, correlations between plasmids and *Wolbachia* sequencing depths were analyzed using the *Wolbachia* DNA library spike-in that was added to the plasmid libraries at an equimolar ratio.

Associations between *w*AlbA, *w*AlbB and plasmid abundance were analysed using Standardized Major Axis (SMA) regression implemented in the smatr R package (Warton et al. 2012). SMA regression was used because both variables in each comparison were estimated from sequencing depth and were therefore subject to measurement error. Sequencing depths were log_2_-transformed after adding a pseudocount of one, and residualized against mitochondrial sequencing depth prior to analysis. The association between *w*AlbA and *w*AlbB was tested using these mitochondrial-depth-corrected residuals. For plasmid analyses, partial SMA regressions were performed by additionally removing the effect of the non-focal *Wolbachia* strain from both the plasmid and focal strain before fitting the final regression.

The effects of genetics, geography and other potential confounding factors on *Wolbachia* density were first analysed using type II ANOVA linear models using the R package car. Since some variables were not fully crossed in our dataset, particularly laboratory-rearing status and population, the full model could not estimate their effects separately. We therefore compared a series of reduced models in which confounded factors were removed in turn. Laboratory-rearing status was not supported as a significant predictor in reduced models where its effect could be estimated, and was therefore excluded from the final model. The final model tested the effects of host mitochondrial subclade, population of origin, mosquito sex and developmental stage while adjusting each term for the remaining variables.

Phylogenetic effects on *Wolbachia* density were tested by fitting generalized linear mixed models using the Bayesian approach implemented in the R package MCMCglmm (Hadfield 2010). All models were ran using flat priors for 13,000,000 iterations with a burn-in of 3,000,000. Model convergence was assessed by visually inspecting the trace of the posterior sample and ensuring the autocorrelation between successive samples in the MCMC chain was below 0.1. Credible intervals were estimated from the posterior distribution of parameter estimates as the 95% highest posterior density intervals (HPD).

The densities of *w*AlbA and *w*AlbB were first analyzed together in a multi-response model with the mitochondrial phylogeny as a random effect. The phylogenetic heritability was calculated as the proportion of total variance explained by the phylogenetic effect estimated by the model. A separate univariate model was run to estimate the phylogenetic heritability of the *w*AlbB-to-*w*AlbA density ratio.

### Phylogenomic analysis

Phylogenomic analysis required additional filtering by depth of coverage, using GATK VariantFiltration to filter by depth ≥ 15 (-G-filter "DP < 15") which was run after the hard filtering steps listed above. Following this, bcftools consensus (Danecek et al. 2021) was used to imprint each sample genotype onto a fasta file of the reference assembly, replacing missing and filtered regions with ‘N’ (-a “N”). Fasta files were concatenated, then converted to nexus files using AliView (Larsson 2014). Sites were partitioned into five genomic subsets: 1^st^ codon positions, 2^nd^ codon positions, 3^rd^ codon positions, non-coding RNA, and non-coding intergenic DNA.

Evolutionary analysis was run in Beast v1.10.4 (Drummond et al. 2012) using an HKY substitution model with a gamma site heterogeneity model, an uncorrelated relaxed clock, a constant size coalescent tree prior (Suchard et al. 2018), and 100,000,000 MCMC runs. Independent substitution models were set for the five partitions. XML files were produced in BEAUTi (Drummond et al. 2012) and edited to include product statistics of the evolutionary rates of each partition (mu * default.clock.rate). A lognormal prior was set on the mtDNA 3^rd^ codon rate with x̄ = -16.59613 and sd = 0.0333; these set the evolutionary rate to be identical to *D. melanogaster* (Richardson et al. 2012). The default clock rate was set using a CTMC prior (Ferreira and Suchard 2008). Trees were annotated using TreeAnnotator (Drummond et al. 2012) and visualised with FigTree v1.4.4 (http://tree.bio.ed.ac.uk/software/figtree/). All runs were analysed in Tracer v1.7.2 (Rambaut et al. 2018).

### *Wolbachia* and plasmid gene content analysis

We estimated the copy number of each gene in each genome by dividing its median sequencing depth by the genome-wide depth. Median sequencing depth per *Wolbachia* gene was calculated from the per-site depth calculated in VCFtools v0.1.16 (Danecek et al. 2011) on samples that had not undergone depth filtering. Since reads mapping to multiple locations or genomes were excluded from our analysis, this caused gene copy numbers to be slightly overestimated. To account for this bias, we first discarded genes where >95% of the samples for a given *Wolbachia* strain had copy number < 0.8, as these were likely genes with sequences that partially or completely matched multiple locations or genomes. From the remaining genes, the average median gene depth per genome was re-calculated and used to recalibrate gene copy numbers. Plasmid gene copy numbers were estimated using the same method.

## Acknowledgments

We thank Ary Hoffmann, Qiong Yang, and Nancy Endersby-Harshman for assisting with samples.

TLS was funded by the Australian Research Council (DE230100257) and by a University of Melbourne Faculty of Science ECR Global Collaboration Award. JM and SPS were funded by the Wellcome Trust (grant numbers 202888/Z/16/Z and 226166/Z/22/Z). FJ and JD were funded by the Biotechnology and Biological Sciences Research Council (BB/V000667) and the Natural Environment Research Council (NE/V011979).

## Supplementary Figures

**Figure S1.**
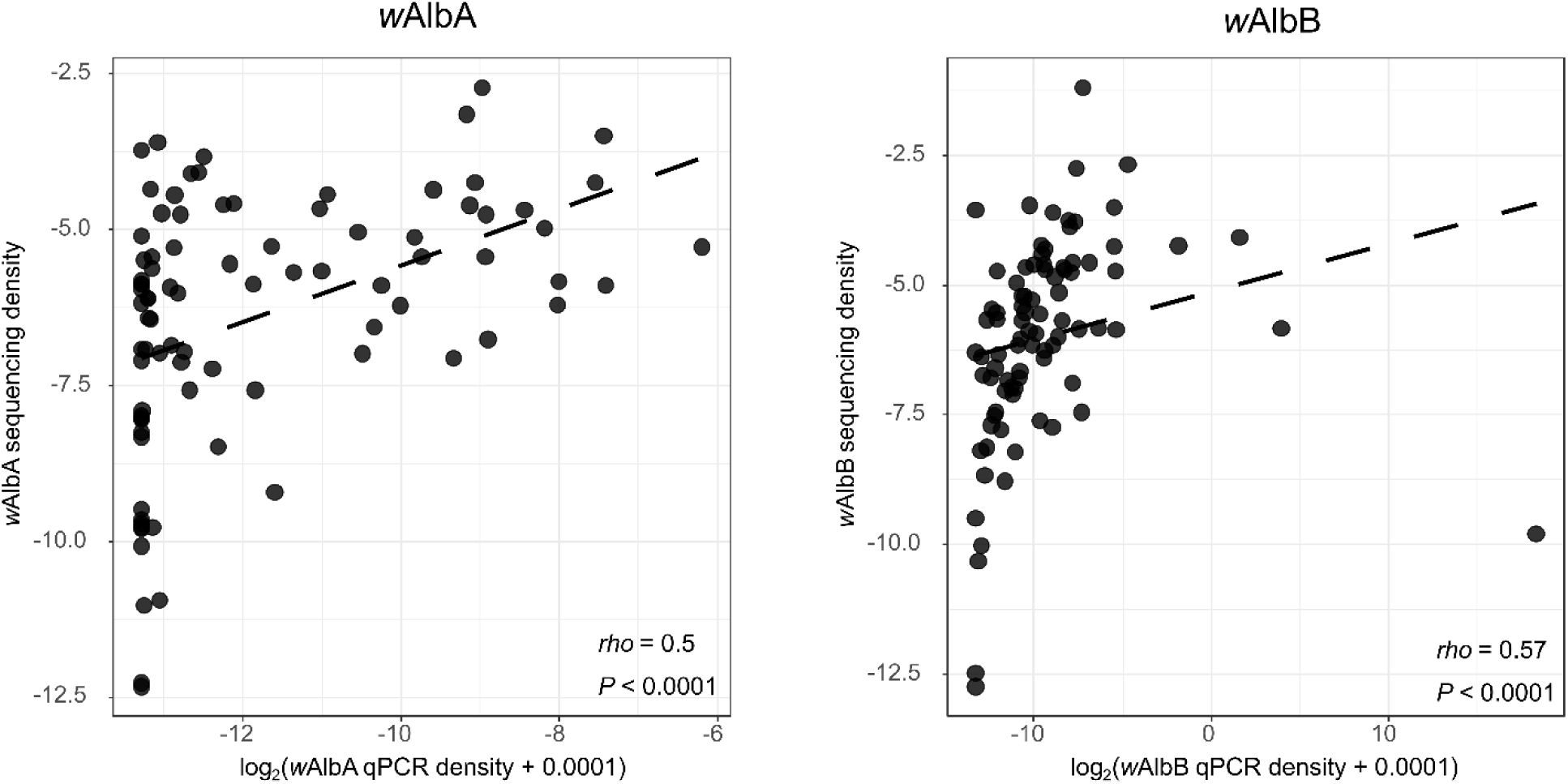
Comparing *Wolbachia* density estimates obtained from qPCR and from sequencing. Correlations between qPCR-based and sequencing-based density estimates are shown for *w*AlbA and *w*AlbB. Correlation statistics correspond to Spearman’s rank correlation tests.

**Figure S2.**
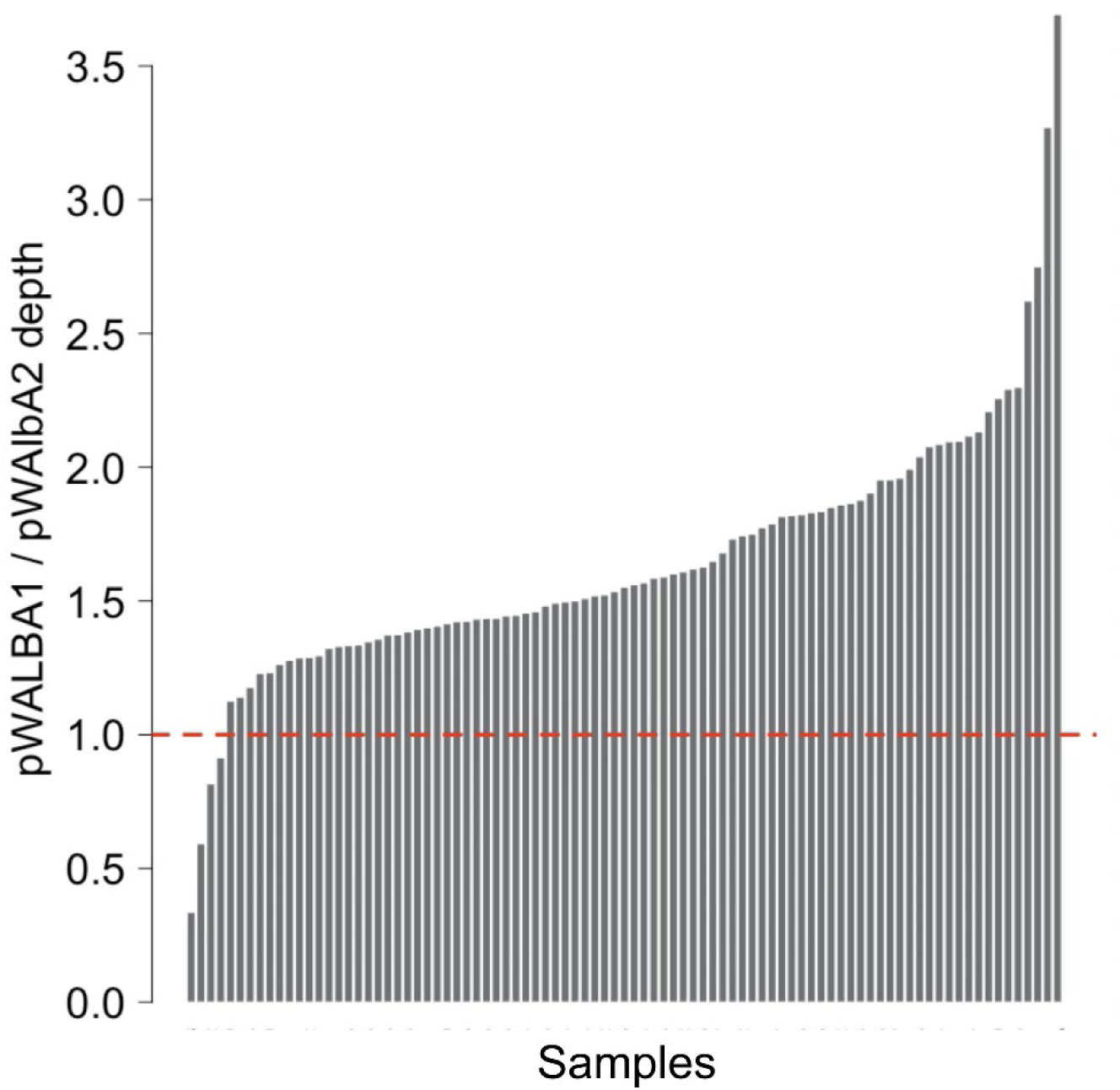
Ratio between pWALBA1 and pWALBA2 sequencing depths per sample.

**Figure S3.**
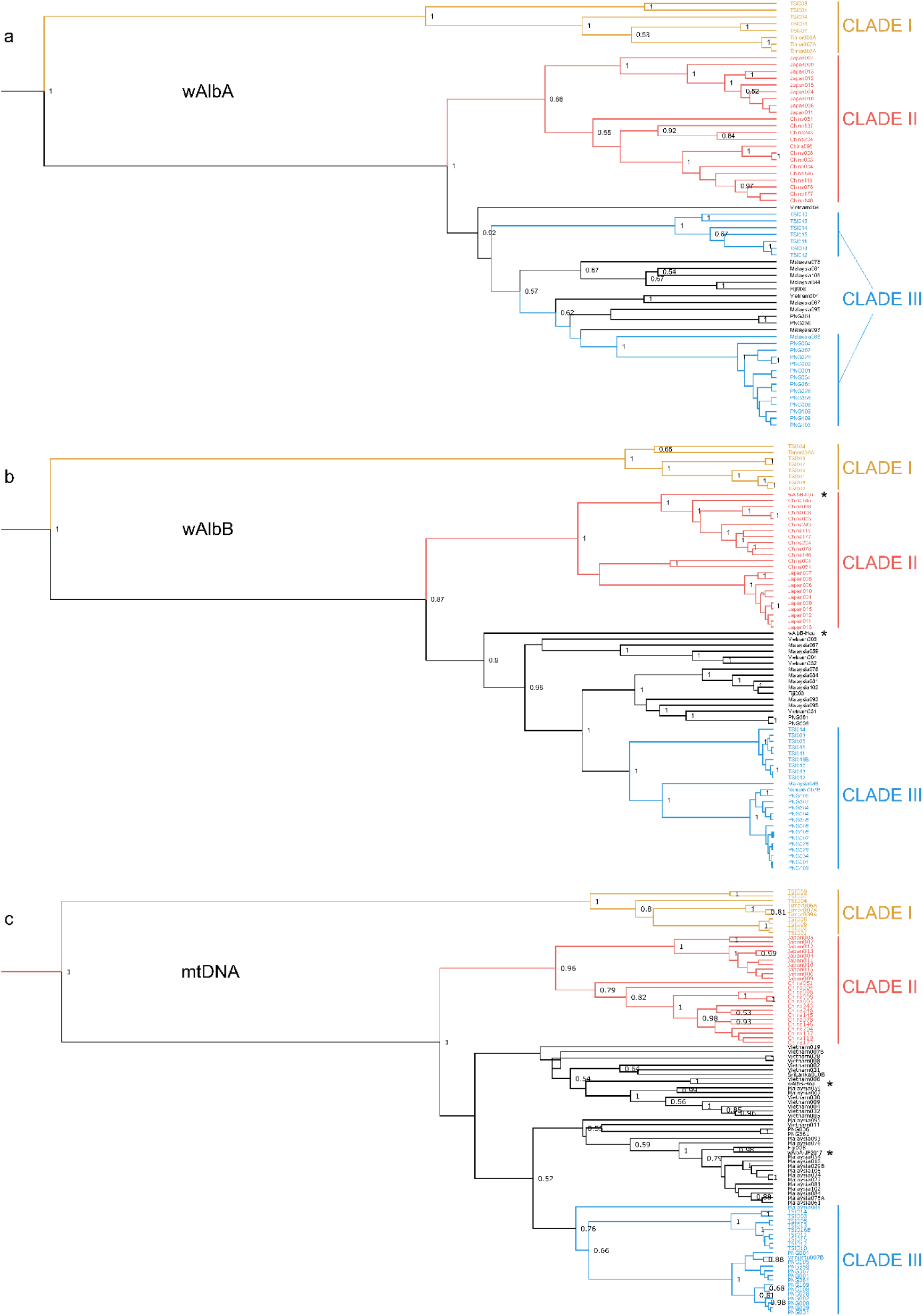
Phylogeny of the *Wolbachia* and mtDNA genomes. Maximum clade consensus trees constructed in Beast v1.10.4 using an HKY substitution model with a gamma site heterogeneity model, an uncorrelated relaxed clock and a constant size coalescent tree prior (Suchard et al. 2018). Independent substitution models were set for 1^st^ codon positions, 2^nd^ codon positions, 3^rd^ codon positions, non-coding RNA, and non-coding intergenic DNA. Posterior support is plotted for nodes with ≥0.5 posterior support. Three major clades described in the main text are indicated by colour. Asterisks indicate previously published genomes: *w*AlbB-Uju, *w*AlbB-Hou, and *w*AlbA-JF2017.

**Figure S4.**
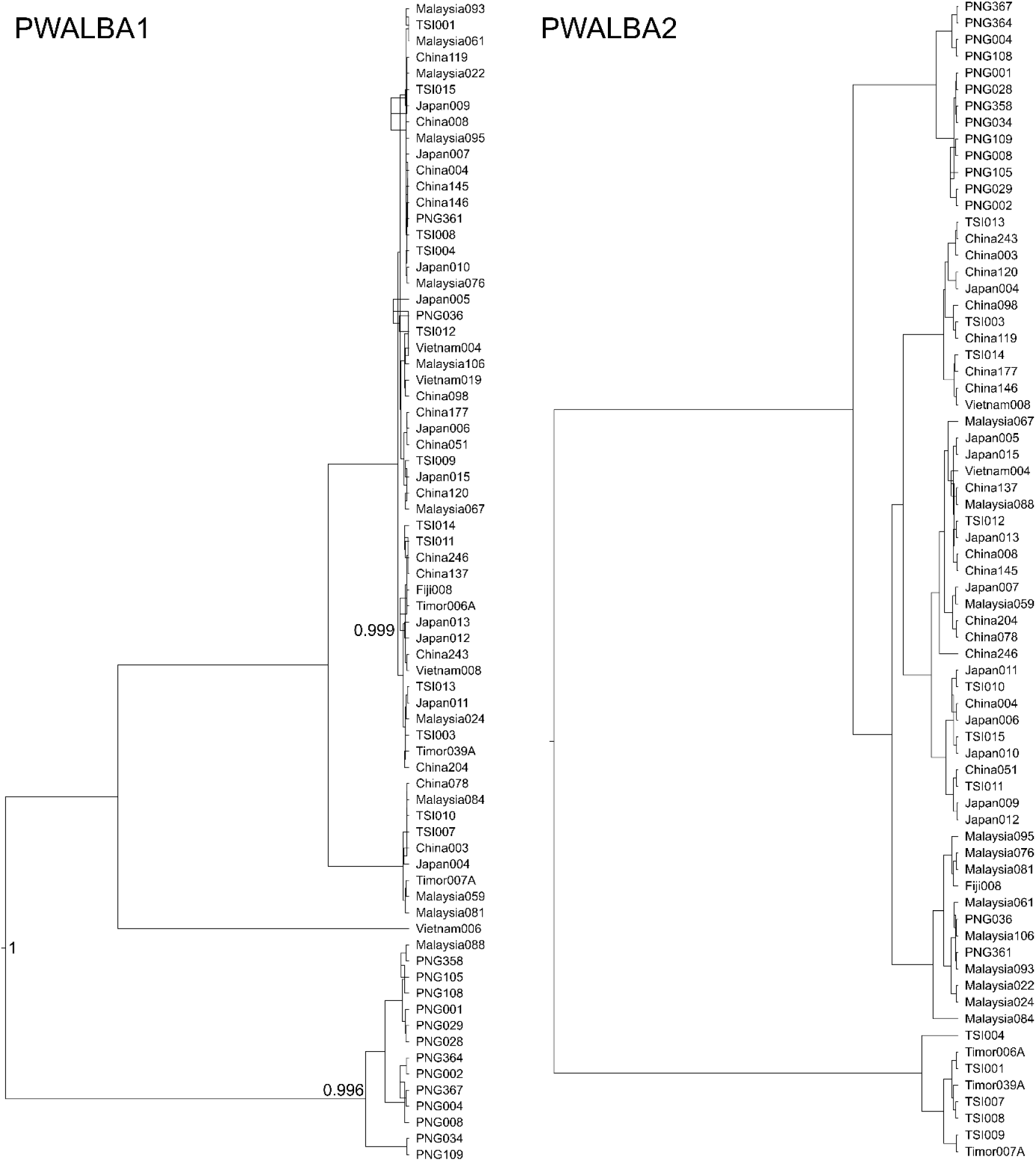
Phylogeny of the two plasmid genomes. Maximum clade consensus trees constructed using Beast v1.10.4 with an HKY substitution model with a gamma site heterogeneity model, an uncorrelated relaxed clock and a constant size coalescent tree prior (Suchard et al. 2018). Independent substitution models were set for 1st codon positions, 2nd codon positions, 3rd codon positions, non-coding RNA, and non-coding intergenic DNA. Numbers at nodes indicate posterior support, plotting only values ≥0.5.

**Figure S5.**
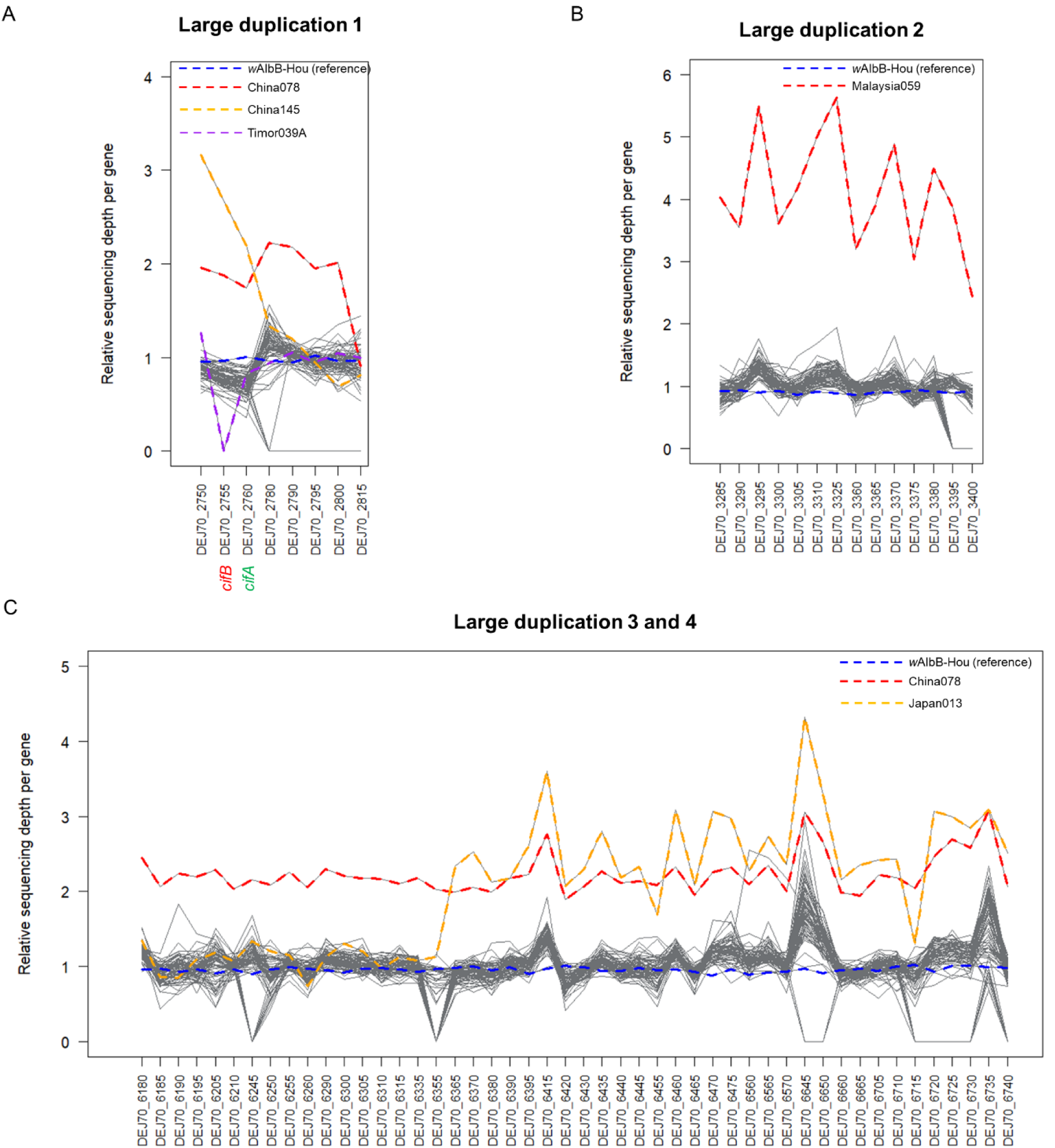
Estimated gene copy numbers in *w*AlbB large duplications per sample and per gene. Each grey line represent a given sample. Samples of interest are coloured with dashed lines for visualization.

**Figure S6.**
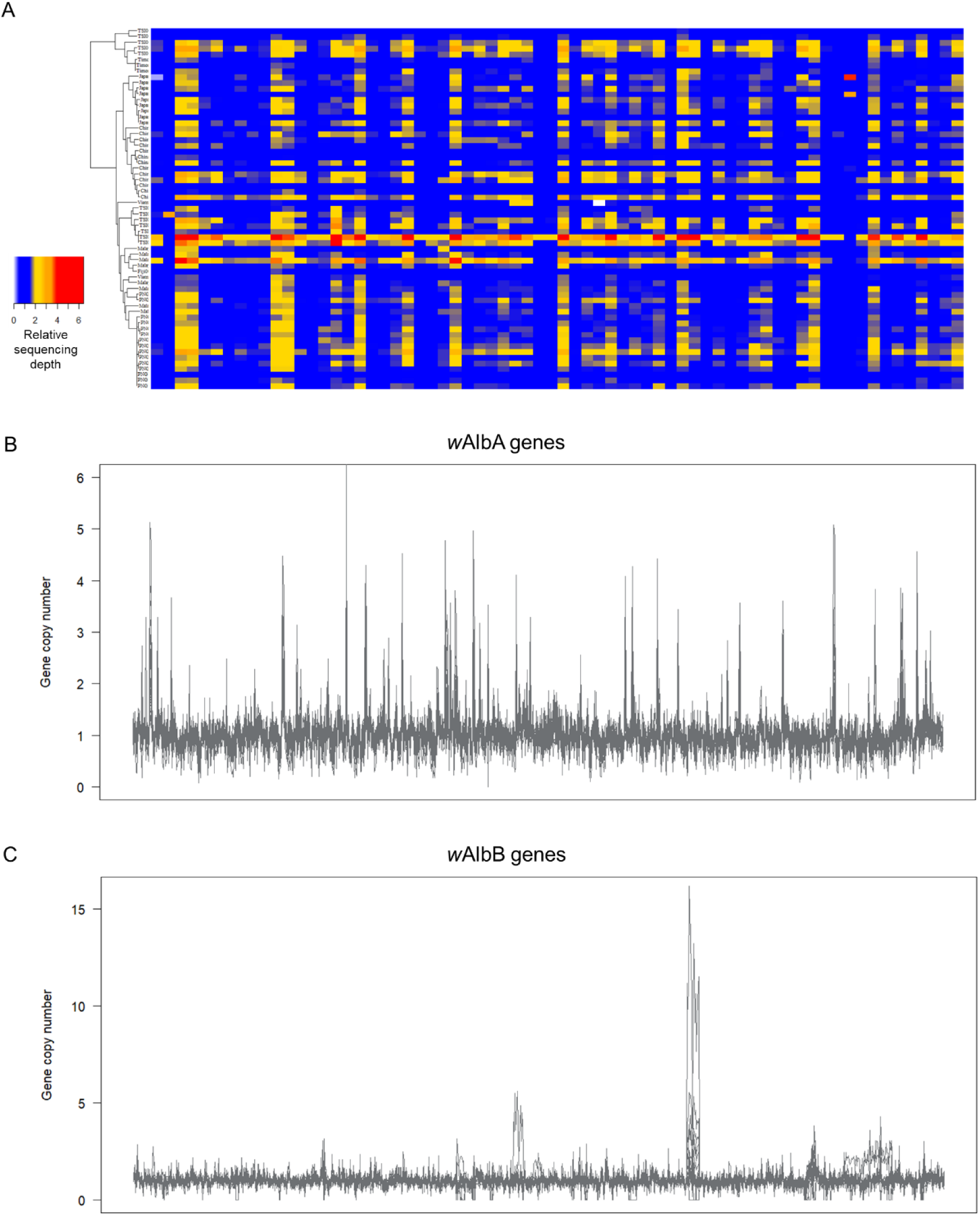
*w*AlbA gene content variation. (A) Left: *w*AlbA phylogeny. Estimated gene copy numbers of *w*AlbA genes (columns) with copy number >1 in at least one sample. (B) *w*AlbA and (C) *w*AlbB gene copy numbers across their respective genomes.

**Figure S7.**
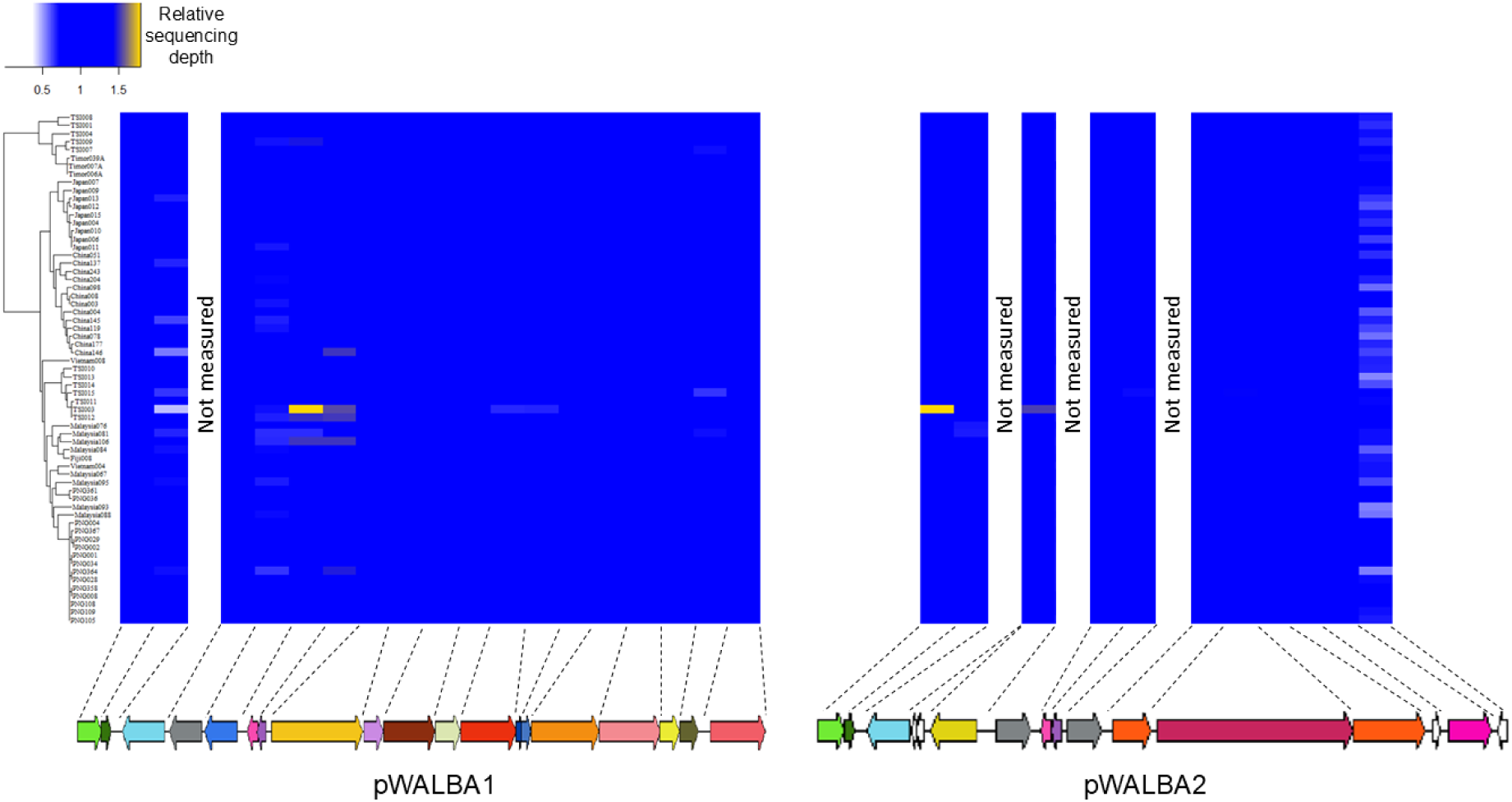
Plasmid gene copy numbers. Left: *w*AlbA phylogeny. Heatmaps: estimated copy numbers of plasmid genes (columns) in each *w*AlbA-infected sample (rows). Bottom: plasmid genome architecture.

